# Reward-specific satiety affects subjective value signals in orbitofrontal cortex during multi-component economic choice

**DOI:** 10.1101/2020.07.04.187518

**Authors:** Alexandre Pastor-Bernier, Arkadiusz Stasiak, Wolfram Schultz

## Abstract

Sensitivity to satiety constitutes a basic requirement for neuronal coding of subjective reward value. Satiety from natural on-going consumption affects reward functions in learning and approach behavior. More specifically, satiety reduces the subjective economic value of individual rewards during choice between options that typically contain multiple reward components. The unconfounded assessment of economic reward value requires tests at choice indifference between two options, which is difficult to achieve with sated rewards. By conceptualizing choices between options with multiple reward components (‘bundles’), Revealed Preference Theory may offer a solution. Despite satiety, choices against an unaltered reference bundle may remain indifferent when the reduced value of a sated bundle reward is compensated by larger amounts of an unsated reward of the same bundle; then the value loss of the sated reward is indicated by the amount of the added unsated reward. Here we show psychophysically titrated choice indifference in monkeys between bundles of differently sated rewards. Neuronal chosen value signals in orbitofrontal cortex (OFC) followed closely the subjective value change within recording periods of individual neurons. A neuronal classifier distinguishing the bundles and predicting choice substantiated the subjective value change. Choice between conventional single rewards confirmed the neuronal changes seen with two-reward bundles. Thus, reward-specific satiety reduces subjective reward value signals in OFC. With satiety being an important factor of subjective reward value, these results extend the notion of subjective economic reward value coding in OFC neurons.

**Significance:** On-going consumption reduces the subjective value of rewards to different degrees depending on their individual properties, a phenomenon referred to as sensory-specific satiety. Such value change should be manifested in economic choices, and neuronal signals for subjective economic reward value should be sensitive to reward-specific satiety. We tested monkeys during choice between two options that each contained two different rewards (‘bundles’); the two rewards were prone to different degrees of satiety. On-going reward consumption affected choices in a way that indicated satiety-induced reward-specific change of subjective economic value. Neuronal responses in the monkey orbitofrontal cortex (OFC) followed the differential reduction of subjective economic value. These results satisfy a crucial requirement for subjective reward value coding in OFC neurons.

## Introduction

There are no specific sensory receptors for rewards, and their value is determined by the needs of individual decision makers. Thus, rewards have subjective value rather than being solely characterized by physical measures such as molecular concentrations, milliliters of juice or units of money. Accordingly, neuronal signals in prime reward structures, such as orbitofrontal cortex (OFC) and dopamine neurons, code reward value on a subjective basis (1–3). One of the key factors determining subjective value is satiety that arises from on-going reward consumption. After drinking a cup of coffee, we may desire a glass of water. We feel sated on coffee while still seeking liquid. Apparently, the coffee has lost more value for us than water. Such value loss is often referred to as sensory-specific satiety (or, more appropriately here, reward-specific satiety) and contrasts with general satiety that refers indiscriminately to all rewards (4, 5). Thus, reward-specific satiety is a key factor of subjective reward value, and any claim towards neuronal subjective reward value coding should include sensitivity to reward-specific satiety.

Reward-specific satiety in humans, monkeys and rodents affects approach behavior, goal-directed behavior, operant responding, learning and pleasantness associated with the specific reward. Lesioning, inactivation, and neural studies demonstrate the involvement of frontal cortex, and in particular OFC, in behavioral changes induced by general and reward-specific satiety. The studies assessed alterations of associative strength, cognitive representations for learning, approach behavior and goal-directed behavior (6–18) but did not address the appreciation and maximization of subjective economic reward value that constitute the central interest of economic decision theory (18–20) and current neuroeconomics research (1, 21, 22, 23, 24). Economic reward value cannot be measured directly but is inferred from observable choice (19–21). Value estimations are made at choice indifference between a test option and a reference option, which renders them immune to unselective general satiety and controls for confounding slope changes of choice functions (25). While choice indifference is possible with milder value differences from reward type, reward delay, reward risk or spontaneous fluctuations (1, 3, 26), it may fail with substantial satiety when animals categorically prefer non-sated alternatives (15–17). By contrast, choice indifference becomes feasible when an added amount of unsated reward can compensate for the value loss of the sated reward. Such tests require reward options with two reward components (‘bundles’). Indeed, all choice options constitute bundles; they are either single rewards with multiple components, like the taste and fluid of a cup of coffee, or contain multiple rewards, like meat and vegetables of a meal we choose.

The rationale of our experiment rests on the tenet that candidate neuronal signals for subjective economic value need to be sensitive to reward-specific satiety. We used bundles whose multiple rewards sated differentially and allowed testing at choice indifference. Using strict concepts of Revealed Preference Theory (19, 27, 28), we had demonstrated that monkeys chose rationally between two-reward bundles by satisfying completeness, transitivity and independence of irrelevant alternatives (24). Using these validations, we now estimated the loss of economic value from reward consumption during on-going task performance. Using two-reward bundle options, instead of single-reward options, we tested choice indifference at specifically set, constant levels. We used multiple, equally preferred indifference points for constructing two-dimensional graphic indifference curves (IC) on which all bundles had by definition the same subjective value. The slopes of these ICs demonstrated subjective value relationships (‘currency’) between two bundle rewards. On-going reward consumption changed the IC slopes in characteristic ways that suggested reward-specific subjective value reduction. During full recording periods of individual OFC neurons, chosen value responses tracked the IC slope changes in a systematic way that suggested a neuronal correlate for reward-specific satiety. These data support and extend the claim of subjective economic reward value coding by OFC neurons.

## Results

### Design

This study assumed that subjective reward value can be inferred from observable economic choice, that altered choice indicates changed subjective value, and that reduction of subjective value with reward consumption reflects satiety. By definition, two options that are chosen with equal probability have the same subjective value (indifference at choice *P* = 0.5 each option). Testing at choice indifference is important, as testing at other positions on the sigmoid choice function cannot exclude confounds from slope changes of the choice function (25). We implemented the following design:

1. A standard economic method estimates subjective economic value at choice indifference against a constant reference reward. As animals may not choose a sated single reward when a non-sated alternative is available (15–17) and thus fail to achieve choice indifference, we studied bundles that each contained the same two rewards on which the animals sated to different degrees. Thus, satiety-induced stronger value reduction of one reward could be compensated by larger quantity of the other, less sated reward to maintain choice indifference. Although the animal sated on one reward much less than on its alternative, it chose each bundle on one half of the trials. The choice involved a single arm movement, and the rewards from the chosen bundle were paid out immediately. Thus, the task contained two discrete, mutually exclusive and collectively exhaustive options. Due to the statistical requirements of neuronal data analysis, we tested stochastic choice in repeated trials rather than single-shot choice.
  (1a) Specifically, the animal chose between a Reference Bundle and a simultaneously presented Variable Bundle. In both bundles, Reward A was blackcurrant juice without or with added monosodium glutamate (MSG), on which the animals sated very little. Reward B was either grape juice, strawberry juice, mango juice, water, apple juice or grape juice with added inosine monophosphate (IMG) on which the animals sated substantially, or peach juice inducing less satiety than blackcurrant juice. The height of two bars within two visual stimuli indicated the quantities of Reward A and B of each bundle on a horizontal touch monitor (higher bar indicated more reward) (Fig. 1*A, B*). In the Reference Bundle, both rewards were set to specific test quantities. In the Variable Bundle, one reward was set to a current test quantity, and the quantity of the other reward was adjustable.
2. Choice indifference indicates equal subjective value between options. At choice indifference against a constant bundle, a change in subjective value of a reward of the alternative bundle needs to be compensated by a change of the other reward of that alternative bundle.
  (2a) Specifically, the compensatory change of Reward A of the Variable Bundle required for choice indifference against the constant Reference Bundle was a measure of subjective value loss of Reward B of the Variable Bundle (Fig. 1*C*). The changed IPs of Variable Bundles relative to the constant Reference Bundle are conveniently graphed on a two-dimensional map (Fig 1*D*). As a common satiety-induced value loss with both bundle rewards might not be detectable as IP change, any IP change may hide non-selective satiety; therefore, an IP change would indicate a value change for a specific reward *relative* to the other bundle reward, rather than an absolute value change of that specific reward.
3. A series of IPs aligns as an IC. Thus, an IP change that indicates subjective value change result in an IC change (Fig. 1*E*).
4. As consequence of changed IPs and ICs indicating subjective value change, the original, physically unchanged bundles to which the animal was indifferent before satiety fail to match the IPs and ICs established during satiety. The mismatch depends on the reward being sated; it is smaller for less sated rewards and larger for more sated rewards.
5. The mismatch between IPs and ICs established before and during satiety represents the major measure of altered subjective reward value by on-going consumption and will be used for describing the altered neuronal coding of subjective reward.

**Fig. 1.**
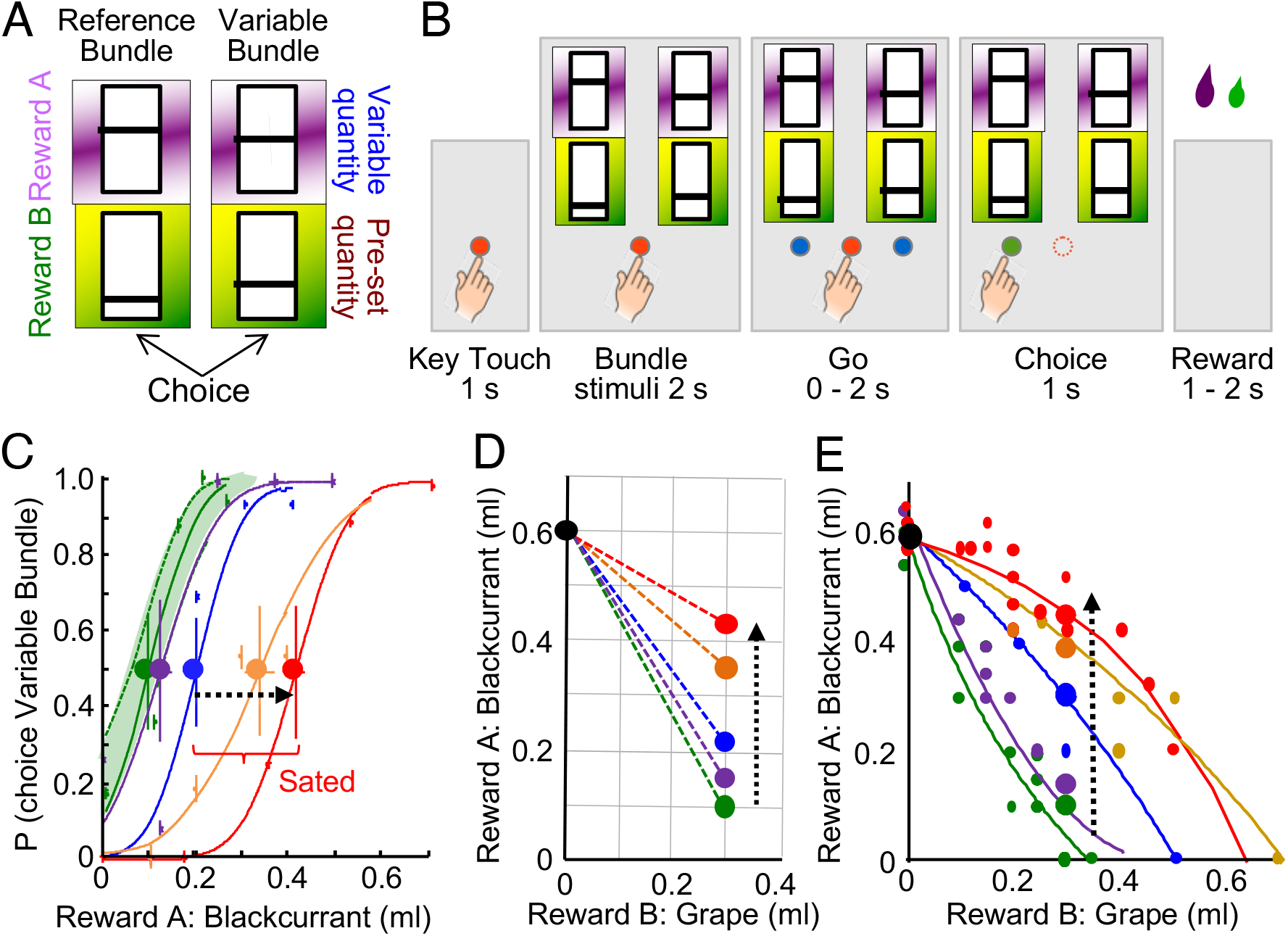
Task, design and psychophysics. (*A*) Choice options (arbitrary reward quantity settings). Each bundle contained two rewards (A, B) with independently set quantities indicated by the vertical bar position within each rectangle (higher was more). The Reference Bundle contained two preset reward quantities. The Variable Bundle contained a specifically set quantity of one reward and an experimentally varied quantity of the other reward. (*B*) Task sequence: In each trial the animal contacted a central touch key for 1.0 s; then the two choice options appeared on a computer monitor. After 2.0 s, two blue spots appeared on the monitor, and the animal touched one option within 2.0 s. After touching the target for 1.0 s, the blue spot underneath the chosen bundle turned green and the blue spot disappeared. The computer-controlled liquid solenoid valve delivered Reward A at 1.0 s after the choice, and Reward B 0.5 s later. (*C*) Psychophysical assessment of choice between constant Reference Bundle (0.6 ml blackcurrant juice, 0.0 ml grape juice) and one of the Variable Bundles (varying reward A, blackcurrant juice, from 0 to 0.7 ml, while holding grape juice constant at 0.3 ml); same bundles as (*C*). Green and violet curves inside green 95% confidence interval: initial choices; blue, orange and red curves: advancing consumption; heavy dots: indifference points (IP). Satiety was defined by IPs exceeding the CI. Each curve and IP were estimated from 80 trials in a single block (Weibull fits, Monkey A. (*D*) Two-dimensional map indicating progressively more blackcurrant juice required for choice indifference between the Reference Bundle and one of the Variable Bundles (from green to red); same test as (*C*). Black and colored dots show bundle positions, connecting dotted lines indicate choice indifference between the connected bundles (IPs). The flattened slope indicated subjective value loss of grape juice relative to blackcurrant juice. (*E*) Gradual changes in slope and curvature of ICs between pre-satiety (green, violet) and during increasing satiety (blue, orange, red). Each IC was estimated from fitting to about 35 IPs (Eq. **1**), with 80 trials/IP (Monkey A). Small dots indicate IPs, large dots indicate IPs estimated from a single psychophysical test sequence (*C, D*).

### Consumption-induced relative subjective value reduction

A typical test started with a fixed Reference Bundle (0.6 ml blackcurrant juice, no grape juice) and a Variable Bundle (variable blackcurrant juice quantity; fixed 0.3 ml grape juice). We titrated Reward A of the Variable Bundle (common currency) to achieve choice indifference between the two bundles. Psychophysical estimations with Weibull-fitted IPs during blocks of 80 trials demonstrated choice indifference with 0.1 ml of blackcurrant juice in the Variable Bundle (Fig. 1*C*, leftmost green IP; *P* = 0.5 choice of each bundle). On a two-dimensional plot of bundle rewards (Fig. 1*D*), the connecting line between two bundles indicated equal subjective value: the Variable Bundle (green dot) had the same subjective value as the Reference Bundle (black dot); the two dots were IPs relative to each other. Thus, at choice indifference, the gain of 0.3 ml of grape juice in the Variable Bundle relative to the Reference Bundle had a common currency value of 0.5 ml of blackcurrant juice.

With on-going consumption of blackcurrant and grape juices, choice indifference required increasing blackcurrant juice, as indicated by successive IPs (Fig. 1*C*, from green via violet, blue and orange to red at *P* = 0.5). Apparently, the gained 0.3 ml of grape juice in the Variable Bundle had lost some of its value for the animal, and only larger blackcurrant juice quantities compensated for that loss. The two IPs within the green 95% confidence interval suggested initially maintained subjective value of grape juice, whereas continuing consumption moved the next IPs outside the CI, indicating progressive reduction of subjective grape juice value. The consumption-induced rightward IP progression (Fig. 1*C*) translated into an upward movement of IPs on a two-dimensional graph (Fig. 1*D*, from green to red), thus flattening the slope between the two IPs. Thus, the additional blackcurrant juice required for choice indifference provided a common currency estimate of the subjective value loss of grape juice. Or, compared to the Reference Bundle, the animal gave up progressively less blackcurrant juice to gain the same quantity of grape juice (0.3 ml) (decreased Marginal Rate of Substitution, MRS, of blackcurrant juice for grape juice). These measures indicated increasing subjective value loss of grape juice relative to blackcurrant juice with on-going consumption.

Wider bundle variations demonstrated consistent consumption-induced subjective value changes (Fig. 1*E*). Starting with a Reference Bundle of 0.6 ml of blackcurrant juice alone, the first IP was obtained by setting the Variable Bundle to coordinates inside the x-y graph. Subsequently, the Reference Bundle was set to the first IP, and another IP was estimated from a newly set Variable Bundle. Repetition of this procedure, in pseudorandomly alternating directions to avoid local distortions (29), resulted in a series of IPs. The curvature of ICs fitted to these IPs (see *Materials and Methods*; Eq. **1**) progressed from convex via near-linear to concave (Fig. 1*E*, green via blue to red). The concavity indicated that the animal required substantial grape juice levels before trading in much blackcurrant juice, possibly reflecting the liquid content of the grape juice, which suggested systematic and advancing subjective value loss of grape juice relative to blackcurrant juice. These slope and curvature changes of two-dimensional IC occurred during recording periods of individual neurons and constituted our test scheme for behavioral and neuronal correlates of reward-specific satiety.

As control, we inverted the test scheme by holding blackcurrant juice constant and varying grape juice psychophysically to obtain choice indifference. Here, instead of the common currency quantity of blackcurrant juice, the increasing quantity of grape juice at choice indifference quantified its subjective value decrease with on-going consumption. When using single-reward bundles with only one but different reward in each bundle having non-zero amount, consumption of both rewards increased IPs, flattened IC slopes and increased blackcurrant:grape juice ratios at IPs; when using two-reward bundles, IC curvature changed from convex to concave (Fig. S1*A-D*). The ICs with bundles containing water in Monkey B showed similar slope flattening and reduced convexity with this test scheme (Fig. S1*E, F*). Thus, the consumption-induced relative subjective value reduction of grape juice or water relative to blackcurrant juice was robust irrespective of test scheme.

### Consistency across bundles

Two rhesus monkeys performed 74,659 choices with one or more of the eight bundle types (Fig. 2). We defined the boundary between pre-sated and sated states by the CI of the initial, left-most choice function between blackcurrant juice and any reward (green in Figs. 1*D*, S1*A* and S1*E*); any IP outside this interval indicated subjective value reduction.

**Fig. 2.**
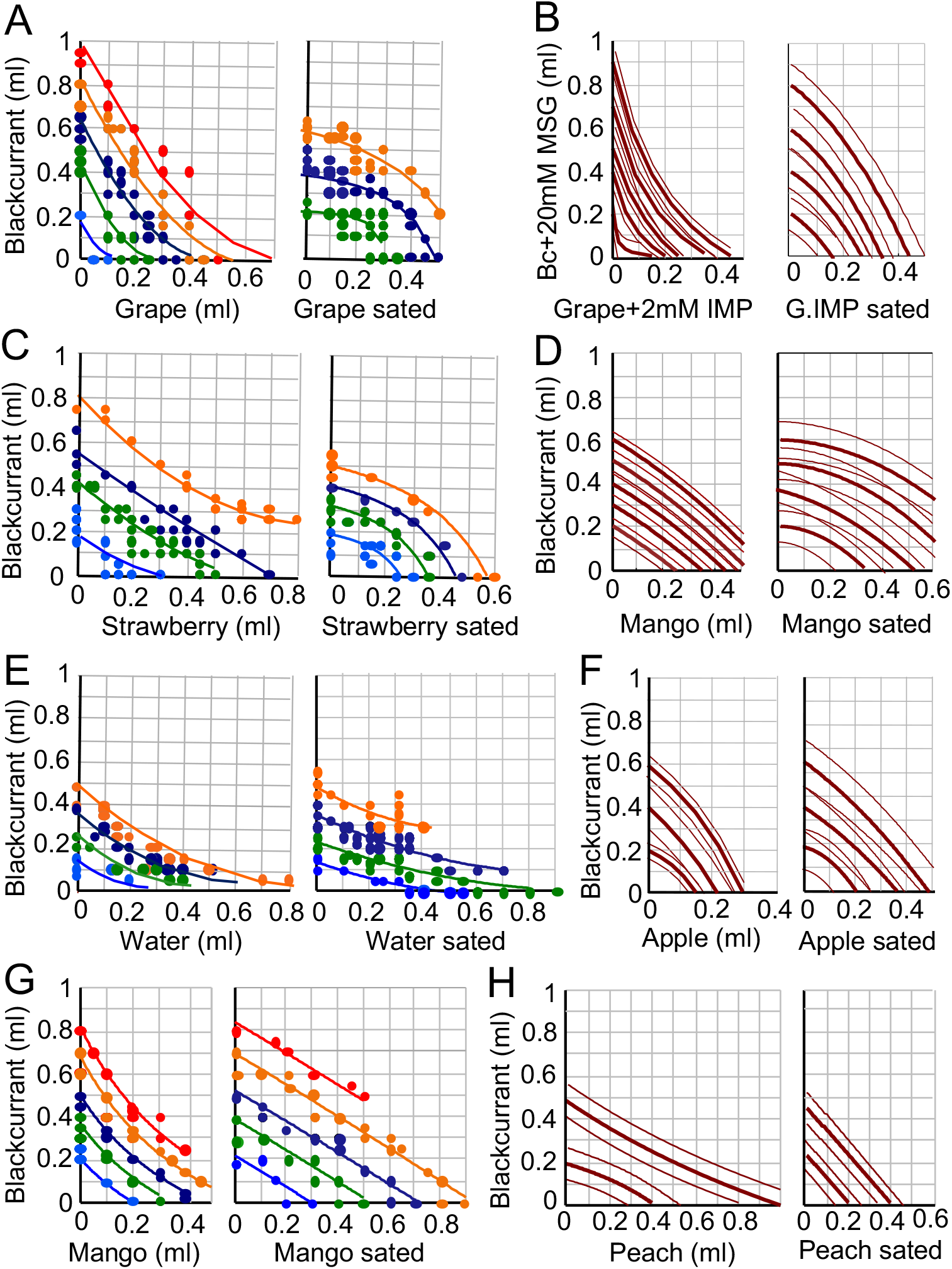
Behavioral effects of reward-specific satiety evidenced by change in choice indifference curves (IC) in all bundles tested. (*A*) - (*F*) Behavioral ICs for all bundle types of the current experiment (Monkey A). Lines show ICs fitted hyperbolically to indifference points (IP) of same color (Eq. **1**). Incomplete lines reflect fitting limited to estimated IPs. Dots in *A, C, E* show measured IPs (choice indifference between all bundles of same color). Thin lines in (*B, D, F, H*) show 95% confidence intervals. Reward A is plotted on the y-axis, Reward B on the x-axis. Bc, blackcurrant juice; MSG, monosodium glutamate; IMP, inosine monophosphate. IMP and MSG were only added to the juices shown in (B). Same color convention in (*A*), (*C*), (*E*) and (*G*) as in Fig. 1*C-E*. Panels (*G*) and (*H*) refer to Monkey B.

Before satiety, we used a total of 38,443 choices to estimate 56 IPs for fitting up to 5 ICs with the bundle (blackcurrant juice, grape juice), 68 IPs for 4 ICs with bundle (blackcurrant juice, strawberry juice) (Monkey A, unless otherwise noted), 58 IPs for 4 ICs with bundle (blackcurrant juice, water), 38 IPs for 5 ICs with bundle (blackcurrant juice, mango juice) (Monkey B), 65 IPs for 5 ICs with bundle (blackcurrant+MSG, grape+IMP), 55 IPs for 5 ICs with bundle (blackcurrant juice, mango juice), 45 IPs for 3 ICs with bundle (blackcurrant juice, apple juice), and 40 IPs for 2 ICs with bundle (blackcurrant juice, peach juice) (Monkey B).

During satiety, we used 36,216 trials to estimate 52 IPs for 3 ICs with bundle (blackcurrant juice, grape juice), 37 IPs for 4 ICs with bundle (blackcurrant juice, strawberry juice), 63 IPs for 4 ICs with bundle (blackcurrant juice, water), 48 IPs for 5 ICs with bundle (blackcurrant juice, mango juice) (Monkey B), 49 IPs for 4 ICs with bundle (blackcurrant+MSG, grape+IMP), 52 IPs for 4 ICs with bundle (blackcurrant juice, mango juice), 55 IPs for 3 ICs with bundle (blackcurrant juice, apple juice), and 44 IPs for 2 ICs with bundle (blackcurrant juice, peach juice) (Monkey B).

On-going reward consumption changed IC curvature from convex to linear and concave or flattened IC slopes, with all bundles tested in this study. The IC change suggested that both animals became more sated across consecutive trials on water, grape juice, strawberry juice, mango juice and apple juice (x-axis) as compared to blackcurrant juice (y-axis) (Figs. *2A-G*; S1*D, F-H*). Nevertheless, the downward slopes of the ICs indicated remaining positive reward value of the sated liquids: increasing amounts of sated liquid (increase on x-axis) required less compensatory unsated blackcurrant juice (decrease on y-axis) for maintaining choice indifference. None of the sated juices had acquired negative reward value, which would have been indicated by an upward IC slope: increasing amounts of negatively valued sated liquid (increase on x-axis) would have required more blackcurrant juice (increase on y-axis) for compensation at IC, which was not the case with the currently used liquids but had been detected with lemon juice, yoghourt and saline (25). The only exception to this pattern of satiety was peach juice: IC steepness increased for this bundle, indicating less satiety than for blackcurrant juice (Fig. 2*H*). These IC changes demonstrate robust and systematic relative subjective value changes with natural, on-going liquid consumption across a variety of bundle types.

### Control for other choice variables

A logistic regression confirmed that bundle choice varied only with the bundle rewards but not with unrelated variables with on-going consumption, such as trial number within block of consecutive trials and spatial choice (Eq. **2**). As before satiety (24), the probability of choosing the Variable Bundle continued to correlate positively with the quantities of its both rewards, and inversely with the quantities of both Reference Bundle rewards (Fig. S1*I*; VA, VB vs. RA, RB). Further, choice probability for the Variable Bundle was anticorrelated with accumulated blackcurrant juice (MA) consumption and positively correlated with grape juice consumption (MB). This asymmetry is explained by the reward quantities at the IPs; as grape juice lost more subjective value than blackcurrant juice during satiety, the animal required, and thus consumed, more grape juice for less blackcurrant juice at the titrated IP. Trial number within individual trial blocks (CT) and spatial choice CL) did not explain the choice. Thus, even with on-going consumption, the animals based their choice on the reward quantities of the bundles and the actually consumed rewards; unrelated variables kept having no significant influence.

### Lick durations

Licking provides a simple measure that could serve as mechanism-independent confirmation for the subjective value changes seen with choices. With on-going reward consumption of both juices, anticipatory licking between bundle stimuli and the first reward decreased gradually with grape juice but changed inconspicuously with blackcurrant juice (Fig. 3*A, B*). This change suggested greater subjective value loss for grape juice than blackcurrant juice. In both animals, cumulative licking varied considerably between the different liquids and decreased significantly during satiety (green) compared to before satiety (pink) (Fig. 3*C-G*). Thus, the licking changes corresponded to the relative reward-specific subjective value changes inferred from bundle choices, IC slopes and curvatures (Fig. 2).

**Fig. 3.**
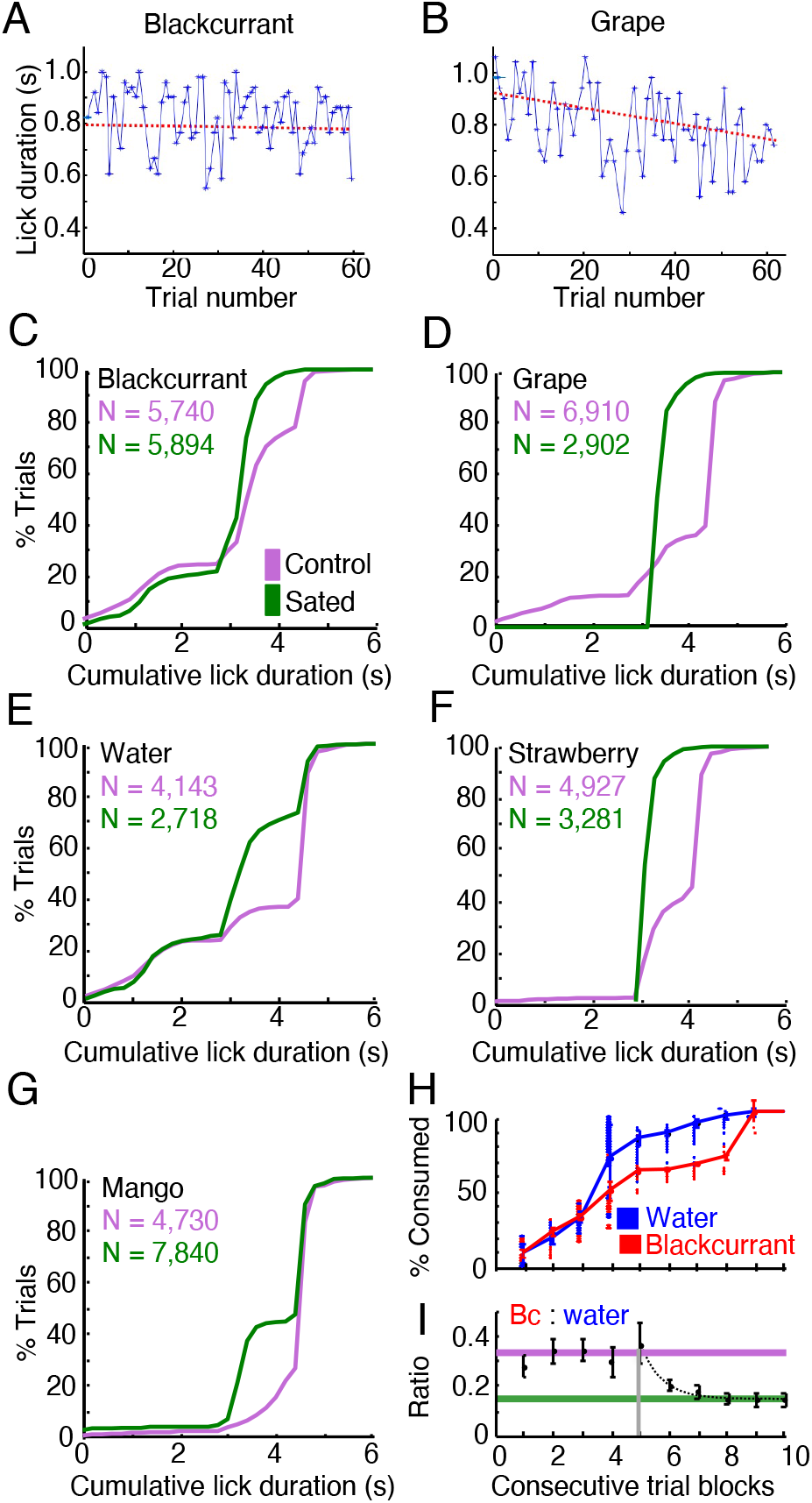
Anticipatory licking and differential juice consumption indicating reward-specific satiety. (*A*), (*B*) Differential decrease of anticipatory licking with single-reward bundles during on-going reward consumption within single test sessions: lick durations remained nearly constant for blackcurrant juice (slope = −2.86 deg, R^2^ = 0.56; 69 trials; linear regression, red), but decreased for grape juice (slope = −20.6 deg, R^2^ = 0.50; 65 trials) (Monkey A). (*C*) - (*G*) Cumulative distributions of lick durations between bundle appearance and reward delivery for several single-reward bundles. Both animals showed significantly more trials with longer lick durations before (pink) than during satiety (green) where animals engaged less frequently in short licks (*D*). Monkey A, blackcurrant juice: *P* = 5.46 x 10^−4^; Kolmogorov-Smirnov test; *n* = 5,740 / 5,894 pre-sated/sated trials) grape juice: *P* = 2.59 x 10^−9^; *n* = 6,910 / 2,902, water: *P* = 3.60 x 10^−3^; *n* = 4,143 / 2,718, strawberry juice: *P* = 8.66 x 10^−6^; *n* = 4,920 / 3,281; Monkey B, mango juice: *P* = 2.41 x 10^−9^; *n* = 4,730 / 7,840. (*H*) Cumulative consumption of water (Reward B) and blackcurrant juice (Reward A) during 10 advancing blocks and 7,160 trials (including bundles with two non-zero quantities). For constant blackcurrant quantities (red), the animal consumed significantly more water than blackcurrant as trials advanced (Monkey A). (*I*) Reduction of blackcurrant:water consumption ratio (Bc:water) from 0.32 (1:3) before satiety (pink) to 0.15 (1:6) with on-going consumption (green). Single exponential function f (β, x) starting at vertical grey line: β_1_ + β_2_e^(β3x)^; [β_1_, β_2_, β_3_] = [0.15, 254.78, −1.41] (β_1_: final ratio, green line; β_2_: decay constant). Consecutive 10 trial blocks for fitting included last block with stable ratio. *n* = 5,520 trials with single-reward bundles; see Fig. S1*A, B* for test scheme); Monkey A.

### Liquid consumption

The IC flattening with on-going consumption indicated that the animal required increasing quantities of the more devalued Reward B for giving up the same quantity of the less devalued blackcurrant juice (Reward A) at IP (Fig. 1*E*). This change was also evident in the choice between the constant Reference Bundle containing only blackcurrant juice (Reward A) and the Variable Bundle containing only one of the other liquids (Reward B) (anchor trials; Fig. S1*B*; increase on x-axis). With on-going consumption, the animal gave up the same quantity of the less sated blackcurrant juice only if it received increasingly more of the sated Reward B at choice indifference. As the animal had no control over the constant Reference Bundle that defined the IP, it ended up consuming more of the devalued reward as the session advanced. For example, with on-going consumption of the bundle (blackcurrant juice, water), consumption increased more rapidly within each day for water (on which the animal was more sated) than for blackcurrant juice (on which the animal was less sated) (Fig. 3*H*; blue vs. red; *P* = 5.0979 x 10^−7^; Kolmogorov-Smirnov test; *n* = 7,160 trials including bundles with two non-zero quantities). The differential consumption change with satiety resulted in decreases of blackcurrant:water consumption ratios at IP (Fig. 3*I*; anchor trials only). These ratio changes indicated relative subjective value changes among the juices (change in common currency).

Similar or less pronounced changes of consumption and consumption ratios occurred with bundles containing blackcurrant juice and grape, mango or apple juice (Fig. S2*A-D*), corresponding to higher satiety for these juices relative to blackcurrant juice (Fig. 2*A, D, F, G*). However, both consumption and ratios increased for bundles combining blackcurrant juice with strawberry or peach juice (Fig. S2*E, F*), corresponding to slightly or noticeably higher satiety for blackcurrant relative to strawberry and peach juice (Fig. 2*C, H*). The ratio changes were consistent across successive weekdays while accelerating on Fridays (Fig. S2*G-J*).

To control for the delivery of two different juices, we tested a bundle containing blackcurrant juice as both components and found no change in consumption ratio (Fig. S3*A*). Further, we reversed the juice sequence, delivering grape juice (Reward A) before blackcurrant juice (Reward B), and interchanged their x-y coordinates. On-going juice consumption consistently changed IC slope and curvature (Fig. S3*B*). The slope became substantially steeper, which corresponded to the flattened slopes with the regular sequence and x-y coordinates (Fig. 2*A*). The curvature became partly concave, indicating that the animal gave up little grape juice for small blackcurrant juice quantities but much more grape juice for larger blackcurrant juice quantities. Together, these IC changes with reversed juice sequence suggested overall reduction of subjective grape juice value relative to blackcurrant juice that confirmed the satiety in the regular juice sequence and indicated grape juice satiety independent of delivery sequence. Further, temporal consumption profiles were similar for both juices (Fig. S3*C* top), whereas consumption ratios reflected well the reduced grape juice value relative to blackcurrant juice (Fig. 3*C* bottom) and corresponded to the inverse ratio changes in the opposite, regular juice sequence (Fig. S2*A*).

Thus, the consumption changes confirmed the relative reward-specific subjective value changes inferred from bundle choices.

### Neuronal test design

Due to the IC change with on-going reward consumption, the original, physically unchanged bundles that were IPs before satiety failed to match the new ICs estimated during satiety. The mismatch depended on the degree of satiety: bundles with variation of less sated reward showed less mismatch, whereas bundles with variation of more sated reward showed more mismatch. To benefit from the IC scheme, we used two tests: variation of blackcurrant juice while holding grape juice constant, and variation of grape juice while holding blackcurrant juice constant. Comparison of IC maps between the pre-sated state (Fig. 4*A* and *B*, left) and the sated state (*C* and *D*, left) shows that IC flattening with satiety moved bundle positions relative to ICs very little for blackcurrant juice variation (*A* vs. *C*) but substantially for grape juice variation (*B* vs. *D*).

**Fig. 4.**
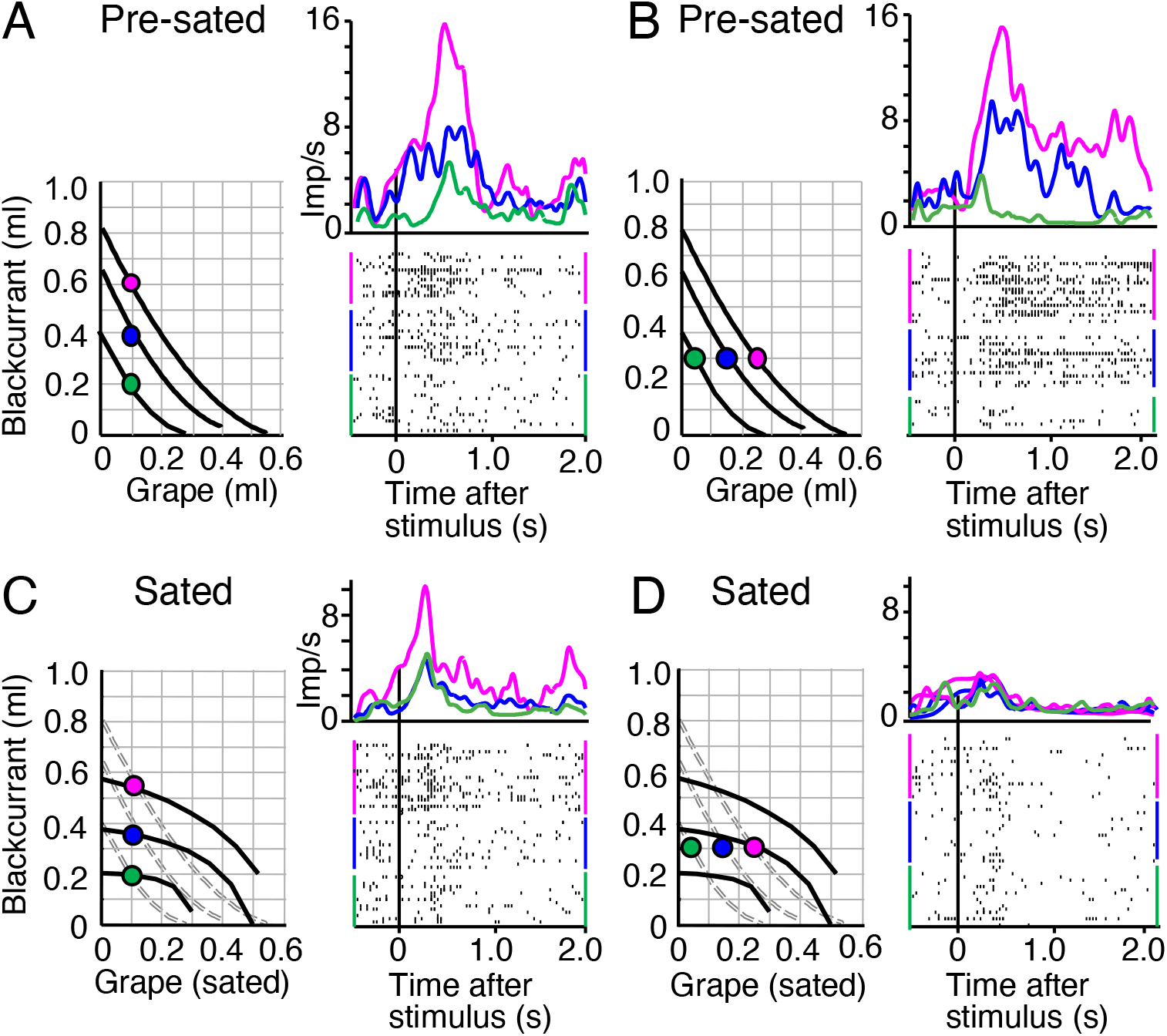
Typical response reduction in single, chosen value coding OFC neuron that follows satiety-related change of choice indifference curves (IC). (A) Monotonic response increase across three ICs with increasing blackcurrant juice before satiety during choice over zero-reward bundle (constant grape juice) (P = 0.030, F(1,35) = 8.86; P = 4.3105 x 10^−7^, F(2,35) = 15.05; two-way Anova: baseline vs. post-stimulus; across 3 bundles). Pre-stimulus activity varied insignificantly across ICs (P = 0.6649, F(2,14) = 0.41; 1-way Anova). Binwidth 10 ms, Gaussian kernel smoothing. Each colored dot indicates a bundle with specific blackcurrant and grape juice quantities located on a specific IC. (B) As (A) but significant response variation with grape juice across ICs (constant blackcurrant juice) (P = 3.15305 x 10^−10^, F(1,35) = 41.57; P = 2.9245 x 10^−24^, F(2,35) = 61.9). Insignificant pre-stimulus variation: P = 0.4273, F(2, 14) = 0.87. Same colors as (A). (C) After consumption of both bundle rewards while recording from same neuron: only mild effect for blackcurrant juice. Despite IC change, the three bundles remained on their three original and separate ICs, and neuronal coding of blackcurrant juice remained significant (P = 0.0275, F(1,35) = 4.88; P = 5.0096 x 10^−28^, F(2,35) = 69.09) but peak response was reduced by 29% (from 15.5 to 11 impulses/s; red) and failed to discriminate between intermediate and low bundles. Insignificant pre-stimulus variation: P = 0.0507, F(2,14) = 3.12. Grey dotted lines indicate ICs before satiety, as in (A). (D) Neuronal response change for sated grape juice: response peak reduction by 75% (from 15.2 to 3.8 imp/s; red), and loss of significant variation (P = 0.0008, F(1,35) = 11.2; P = 0.8053, F(2,35) = 0.22). Insignificant pre-stimulus variation: P = 0.9686, F(2,14) = 0.03. After the consumption-induced slope and curvature change of the ICs (from convex to concave), the three physically unchanged bundles were now on or close to the same, intermediate IC, indicating similar subjective value among them and reflecting satiety for grape juice. Dotted ICs are from pre-sated state. Thus, while continuing to code reward value (C), the responses followed the satiety-induced IC change (D). This reduction of bundle stimulus response was seen in 30 neurons (Table S1).

We tested the influence of on-going reward consumption during the recording period of individual neurons, which allowed us to compare responses of the same neuron between non-sated vs. sated states, as defined by IPs inside vs. outside 95% confidence intervals, respectively (Fig. 1*C*, green zone). As these tests required several tens of minutes with each neuron, neurons not coding chosen value were not further investigated. All satiety-tested neuronal responses followed the basic scheme of ICs: monotonic increase with bundles placed on different ICs (testing bundles with different subjective value), and insignificant response variation with bundles positioned along same ICs (testing equally preferred bundles with equal subjective value) (24). We assessed these characteristics with a combination of multiple linear regression (Eq. **3**), Spearman rank-correlation, and two-way Anova (see *Materials and Methods*). All tested responses belonged to the subgroup of previously studied OFC neurons (30) that were sensitive to multiple rewards and coded the value of the bundle the animal chose (‘chosen value’, as defined by Eqs. 4 and 5). To reduce visual confounds between the two choice options, we used similar visual stimuli that were not identifiable as distinct objects; therefore, we could not test object value or offer value coding as the other major OFC response category (1).

We subjected most neurons to two bundle tests: (i) choice over zero-reward bundle; both rewards were set to zero in one bundle, and the animal unfailingly chose the alternative, non-zero bundle; (ii) choice between two non-zero bundles; at least one reward was set to non-zero in both bundles, and the animal chose either bundle (Table 1). In addition, for comparison with previous studies on single-reward options, we tested all neurons with single-reward bundles in which only one reward was set to non-zero; usually the non-zero reward differed between the two bundles.

### Single-neuron subjective value coding follows IC changes

At the beginning of daily testing, neuronal responses during choice over zero-reward bundle followed monotonically the increase of both bundle rewards (blackcurrant and grape juice), confirming value coding. Stimuli for bundles on higher ICs elicited significantly larger responses (Fig. 4*A, B*) (pre-stimulus increases due to the short inter-trial interval varied insignificantly and were unrelated to trial sequence; Fig. S4). With on-going consumption of both bundle rewards, increasing blackcurrant juice, on which the animal was less sated, remained positioned on different ICs. Correspondingly, neuronal responses continued to vary across ICs (here primarily between the top two ICs) (Fig. 4*C*; red vs. blue-green). By contrast, increasing grape juice did not require much compensation by decrease of blackcurrant juice for maintaining choice indifference, as shown by the flattened and concave ICs (Fig. 4*D* left). The IC change indicated that more grape juice had only little more reward value to the animal. Accordingly, the neuron recorded during this test showed only a small and unmodulated response despite grape juice increase (Fig. 4*D* right; *P* = 0.22; across-bundle factor of two-way Anova); the response peak for the largest grape juice quantity dropped by 75%. Thus, on-going consumption of both juices reduced the subjective value variation of grape juice, and the neuronal responses reflected the reduced value variation, without change in coding rules or reward selectivity.

Similar consumption-induced neuronal changes occurred in choice between two non-zero bundles (Fig. S5). As bundles varying only in blackcurrant juice continued to occupy different ICs, OFC responses continued to increase with blackcurrant juice (Fig. S5*A, C*). By contrast, as original bundles varying only in grape juice were now positioned on lower and fewer ICs, neuronal responses decreased and became less differential (Fig. S5*B, D*; red, blue, green). Responses to the physically unchanged bundle whose position had changed from intermediate to highest IC (hollow blue) now dominated all other responses (Fig. S5*D* right, dotted blue line). Finally, before satiety the bundle containing only 0.6 ml blackcurrant juice had similar subjective value as the bundle with only 0.4 ml grape juice (Fig. S5*B*; hollow and solid blue dots on same IC), and correspondingly drew similar neuronal responses (dotted and solid blue lines), whereas with satiety the same two bundles were positioned on different ICs (Fig. S5*D*; hollow vs. solid blue dot) and correspondingly elicited different responses (dotted vs. solid blue line). Thus, the neuronal responses changed with on-going reward consumption irrespective of choice over zero-reward bundle or true choice between two non-zero bundles.

Thus, OFC neurons continued to code reward value with on-going reward consumption. The responses continued to discriminate well the quantity of blackcurrant juice whose subjective value had changed relatively less (Figs. 4*A* vs. 4*C*, S5*A* vs. S5*C*) but were reduced for grape juice whose value had dropped more, despite unchanged physical grape juice quantities (Figs. 4*B* vs. 4*D*, S5*B* vs. S5*D*). In following the altered ICs, these OFC signals reflected reward-specific relative subjective value changes induced by on-going consumption.

### Neuronal population

Out of 424 tested OFC neurons, 272 showed changes during task performance and were investigated during on-going reward consumption neurons. These neurons were located in area 13 at 30-38 mm anterior to the interaural line and lateral 0-19 mm from the midline; they were parts of the population reported previously (30). Responses in 98 of these 272 task-related OFC neurons (36%) coded chosen value (defined by Eqs. 4 and 5) and followed the IC scheme in any of the four task epochs (Bundle stimulus, Go, Choice or Reward) during choice over zero-reward bundle or choice between two non-zero bundles (Table 1). Of the 98 chosen value neurons, 82 showed satiety-related changes with bundles composed of blackcurrant juice (Reward A) and grape juice, water or mango juice (Reward B) (Tables 2, S1).

Using the scheme of differential consumption-induced IC changes of Figs. 4 and S5, we found that averages of z-scored positive subjective value coding responses in 31 neurons (Table 2) continued to vary with blackcurrant juice quantity (despite peak reduction) (Reward A; Fig. 5*A, B*), whereas responses became insignificant with grape juice, water or mango juice in the same neurons (43% peak reduction) (Reward B; Fig. 5*C, D*). These contrasting, consumption-related neuronal response variations occurred individually in all four task epochs (Table S1).

**Fig. 5.**
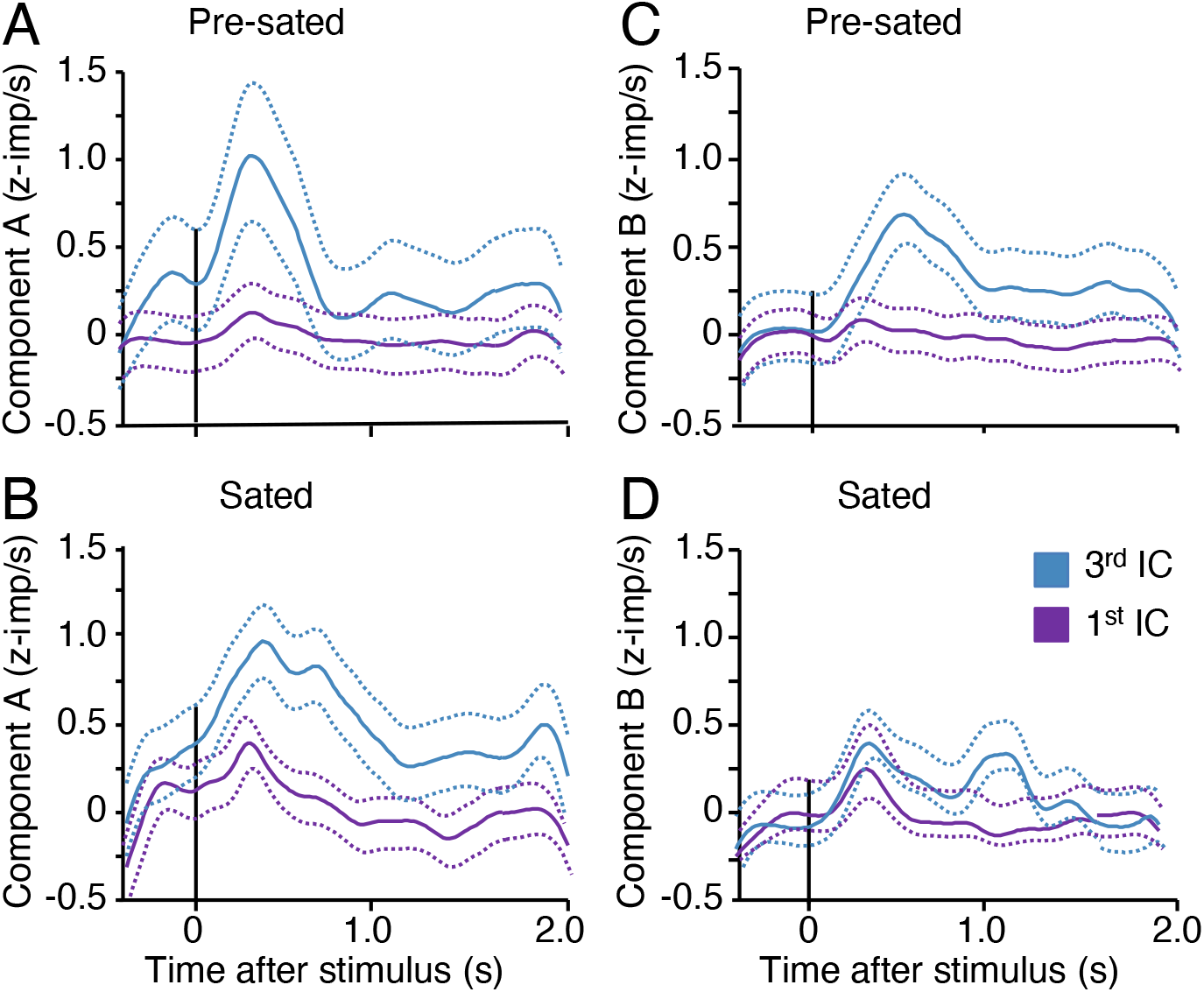
Reduction of neuronal population response with reward-specific satiety. (*A*) - (*D*) Differential satiety-related reduction of averaged z-scored chosen value responses in 30 positive coding neurons from both animals. Each panel shows responses and their 95% confidence interval to bundles on lowest and third lowest indifference curves (IC) during choice over zero-reward bundle. For Reward A (blackcurrant juice), responses differed significantly between lowest and third lowest ICs both before satiety (A: *P* = 0.0015, F(1,3650) = 10.12; *P* = 5.0564 x 10^−4^, F(1,3650) = 19.7; two-way Anova: baseline vs. post-stimulus; across 2 bundles) and during satiety (B: *P* = 3.10864 x 10^−5^, F(1,3853) = 17.39; *P* = 1.35252 x 10^−32^, F(1,3853) = 143.99). By contrast, for Reward B (grape juice, water or mango juice), responses differed significantly before satiety (C: *P* = 0.0005, F(1,5769) = 12.22; *P* = 9.54232 x 10^−10^, F(1,5769) = 38.97), but not during satiety (D: *P* = 0.0028, F(1,4790) = 9.65; *P* = 0.91, F(1,4790) = 0.01).

Quantifications of individual response changes demonstrated consumption-induced significant response reduction with positive subjective value coding neurons and significant response increase with negative (inverse) coding neurons (lower response with higher subjective value) during choice over zero-reward bundle (Fig. S6*A, B*, red) and during choice between two non-zero bundles (Fig. S6*C, D*, red; Tables 2, S1). Fewer neurons showed inverse changes that were difficult to reconcile with subjective value coding (black in Fig. S6), or insignificant changes.

Thus, OFC neuronal population responses reflected consumption-induced changes in subjective value in a similar way as individual neurons.

### Neuronal classifier performance

We used a neuronal classifier as additional means for demonstrating changes of neuronal reward coding by on-going reward consumption. We analyzed only neuronal responses that followed the IC scheme; they increased or decreased monotonically across ICs during any of the four task epochs but changed only insignificantly along ICs; further, the responses changed with on-going reward consumption as shown in Figs. 4, 5 and S5.

We first established the accuracy with which an ideal observer using neuronal responses distinguished bundles on different ICs. We trained a support vector machine (SVM) classifier on neuronal responses to randomly selected bundles positioned on the lowest and highest of three ICs (Bundle stimulus epoch). Before satiety, as defined by IPs inside the initial CI (Fig. 1*C*), the classifier showed decent bundle discrimination with as few as five neurons during choice over zero-reward bundle; classifier performance increased meaningfully with added neurons (Fig. 6*A* black). Then we used this classifier trained on pre-satiety responses and tested distinction of the same bundles using neuronal responses during consumption-induced satiety. We found a significant difference of accuracy (F(1, 34) = 68.02, p = 1.26653 x 10^−9^; 1-way Anova); the accuracy increase with added neurons suggested maintained valid classification (Fig. 6*A* red). The satiety-related accuracy drop was also evident in the inverse test sequence: whereas accuracy was high with classifier training and testing during satiety, it was significantly lower when training on neuronal responses during satiety but testing using responses before satiety (F(1, 34) = 17.99, p = 0,0002).

**Fig. 6.**
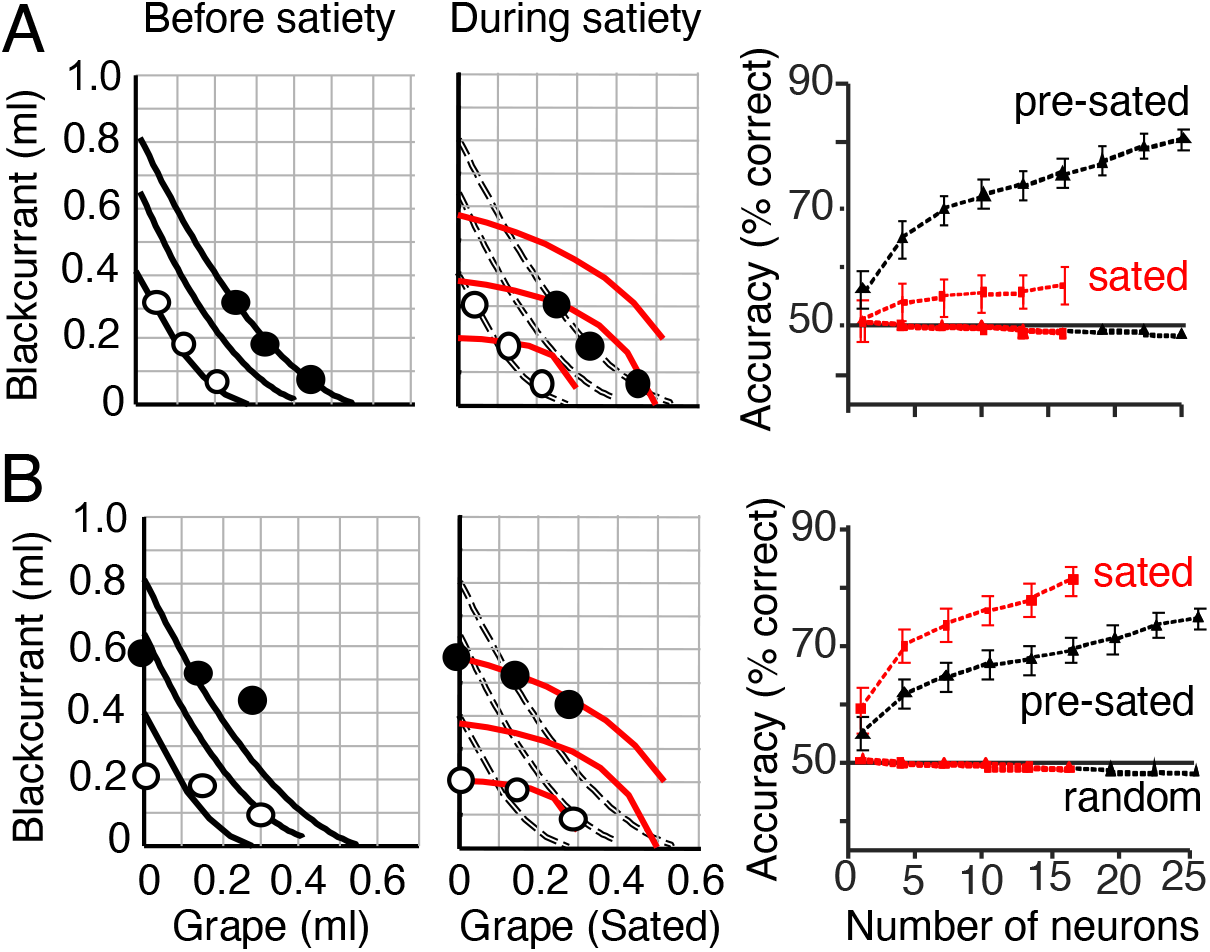
Bundle classification demonstrates satiety-related change of neuronal subjective value coding. (*A*) Bundle classification by support vector machine using neuronal responses to stimuli of bundles positioned on the lowest and third lowest indifference curve, respectively (choice over zero-reward bundle; Bundle stimulus epoch). The classifier was trained on neuronal responses collected before satiety and tested for bundle distinction before satiety (black) and during satiety (red). Left: identical bundle positions on two-dimensional map but IC change with on-going consumption, indicating satiety-induced relative subjective value change (red). Right: classifier accuracy increase with neuron numbers before satiety (black), but drop when tested during satiety (red). Error bars indicate standard errors of the mean (SEM). Random: control classification with shuffled assignment of neuronal responses to tested ICs. (*B*) As (*A*) but reverse testing order: classifier trained on neuronal responses collected during satiety and tested during satiety (red) and before satiety (black).

The change in classifier accuracy occurred with choice over zero-reward bundle with neuronal responses to Bundle stimuli (Fig. 6) and during the Go epoch (Fig. S7*A*), but not during Choice and Reward epochs (Fig. S7*B, C*). The changes were not explained by baseline changes during the 1 s Pretrial control epoch (Fig. S7*D*). Similar accuracy differences were seen in choice between two non-zero bundles during Bundle stimuli, Go epoch and Choice epoch, but not during the Reward epoch (Fig. S7*E-H*), again unexplained by baseline changes (Fig. S7*I*). The accuracy differences were consistent across on-going consumption steps (Fig. S7*J*).

Finally, we analyzed activity from 265 unmodulated and unselected neurons, as before (27). From the total of 424 tested OFC neurons, we excluded the 98 neurons whose activity followed the IC scheme and further 61 neurons that coded only one of the two bundle rewards. Of the remaining 265 neurons, 113 showed task-related activity, whereas the other 152 neurons did not. The classification from the activities of the 265 neurons still showed mild differences between pre-satiety training responses and satiety test responses during the task epochs of Bundle stimuli, Go and Choice (Figs. S8, S9). These remaining differences may reflect sub-significance changes and suggest that subjective value coding reflects satiety mildly in a rather large OFC population.

Thus, the classifier results provide additional evidence for neuronal responses reflecting consumption-induced subjective value changes indicative of satiety.

### Neuronal changes with single-reward bundles

Our use of choice options with two reward components differs from the conventional use of single reward options (1, 2) and thus requires controls and additional analyses. To do so, we used the same two visual component stimuli but set only one, but different, reward in each bundle to a non-zero quantity. The reward settings allowing choice indifference were derived from our use of two-reward bundles. The single-reward bundles were positioned graphically along the x-axis and y-axis but not inside the IC map (Fig. S1*B*) and thus were equivalent to single-reward choice options tested earlier (1, 3, 26). Trials with these anchor bundles interleaved with trials using two regular bundles of which at least one contained two non-zero rewards. These regular bundles provided the IPs at the axes required for analyzing anchor bundle choices.

First, we confirmed the results from two-reward bundles using the same IC formalism. We tested single-reward bundles on the same 82 neurons that had shown satiety-related changes with two-reward bundles. The neuronal responses of Fig. 7*A, B* distinguished both blackcurrant juice and water quantities during choice over zero-reward bundle before satiety. On-going consumption of both rewards flattened the ICs. The IC relationship of blackcurrant juice was preserved, and the neuron kept discriminating blackcurrant juice quantities (Fig. 7*C*). By contrast, the large water quantity was now positioned further below the top IC than before (Fig. 7*D*, red on x-axis) and close to the IC of the small blackcurrant quantity (blue on y-axis), and the small water quantity was now positioned below its original IC (blue on x-axis). Correspondingly, neuronal activity with large water quantity lost its peak and varied only weakly between the two water quantities (Fig. 7*D*, blue vs. red). These neuronal changes with single-reward bundles reflected the consumption-induced subjective value changes in a similar way as with two-reward bundles (Figs. 4, S5).

**Fig. 7.**
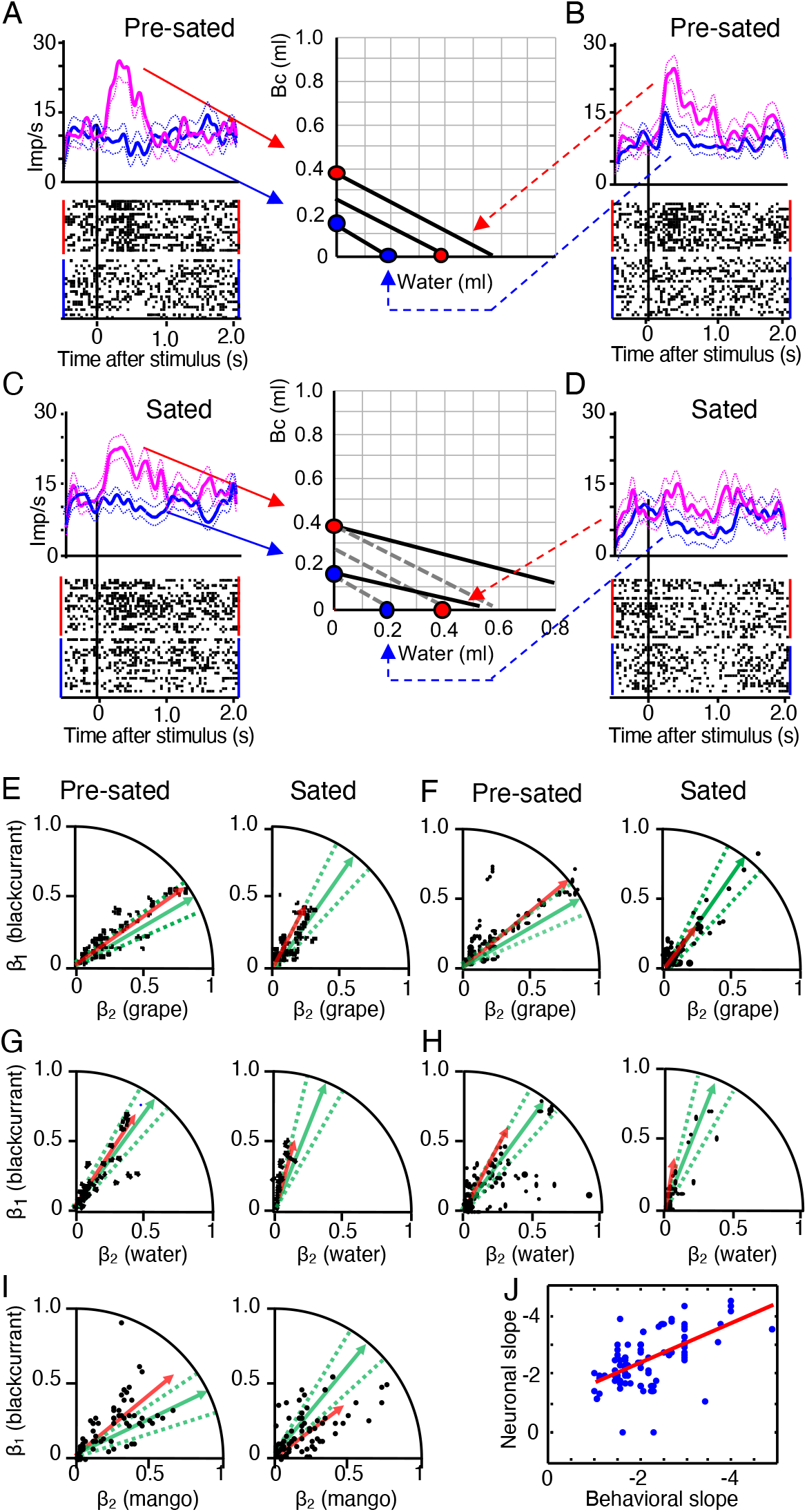
Reward-specific satiety confirmed with single-reward bundles. (*A*) - (*D*) Responses of one chosen value coding neuron before and during satiety. Each bundle contained specific non-zero quantities of only blackcurrant juice or only water (colored dots on x- and y-axes) and was tested during choice over zero-reward bundle. Indifference curves (IC) were obtained from indifference points estimated in interleaved trials with two-reward bundles. Binwidth 10 ms, Gaussian kernel smoothing, 95% confidence interval. (*A*) Significant response increase across two ICs with increasing blackcurrant juice (Bc) before satiety (water remained zero) (*P* = 9.54232 x 10^−10^, F(1,40) = 38.97; *P* = 8.3368 x 10^−10^, F(1,40) = 39.27; two-way Anova: baseline vs. post-stimulus; red vs. blue bundle). (*B*) As (*A*) but significant response variation with increasing water across two ICs (blackcurrant juice remained zero) (*P* = 9.8258 x 10^−6^, F(1,40) = 20.02; *P* = 2.0472 x 10^−8^, F(1,40) = 32.54). Same colors as (*A*). (*C*) Despite IC flattening with on-going reward consumption, the two bundles with blackcurrant juice variation remained on the same two ICs, and the neuronal response variation remained significant (*P* = 4.7616 x 10^−8^, F(1,40) = 30.91; *P* = 6.9739 x 10^−13^, F(1,40) = 54.5), with some peak response reduction (red). Dotted ICs are from pre-sated state. (*D*) IC flattening with reward consumption indicates relative subjective value reduction of water. The two unchanged bundles with water variation were now located on and below the lower IC (dotted lines). Neuronal activity varied only weakly (red) (*P* = 8.9470 x 10^−8^, F(1,40) = 29.6; *P* = 0.0367, F(1,40) = 4.39). Further, the large-water bundle (dotted red line) elicited now a similar response as the low-blackcurrant bundle that was now on the same IC (solid blue line in (*C*)). Thus, while continuing to code subjective reward value (shown in (*C*)), the responses followed the satiety-induced IC change. This reduction of bundle stimulus response was seen in 30 neurons (Table S1). (*E*) Vector plots for behavioral choice of bundle (blackcurrant juice, grape juice) over zero-reward bundle (green) and corresponding z-scored neuronal population responses (rectified for inverse coding) of all 32 tested chosen value neurons, unselected for satiety change (black, red). Neuronal vector slopes were 35 deg before satiety and 62 deg during satiety, pooling all significantly positive and normalized negative (inverse) coding responses from all four task epochs (Tables 1 and 2); all included responses followed the IC scheme. Dots refer to neuronal responses, vectors represent averages from behavioral choices (green; dotted lines: 95% confidence interval) and neuronal responses (red), based on Eqs. 1a and 3, respectively (see *Materials and Methods*). Neuronal slope regression coefficients (β’s) on axes refer to Eq. **3**. (*F*) As (*E*) but for choice between two non-zero bundles. Neuronal vector slopes were 38 deg before and 45 deg during satiety. (*G*), (*H*) As (*E*, *F*) but for bundle (blackcurrant juice, water) (*N* = 33 unselected chosen value neurons). (*I*) As (*E*) but for bundle (blackcurrant juice, mango juice) (*N* = 21 unselected chosen value neurons). The larger deviation between behavioral (green) and neuronal (red) vectors for mango juice (*I*) compared to grape juice and water (*E*) - (*H*) may be due to fewer neurons tested and fewer tests (Table 1). (*J*) Correlation between rectified neuronal and behavioral IC slopes (β’s from Eqs. 3 and 1a, respectively) during satiety in all tested neurons (rho = 0.604; *P* = 8 × 10−6, Pearson correlation; rho = 0.595, *P* = 2 × 10−5, Spearman rank-correlation; *n* = 90 responses during choice between two non-zero bundles).

Next, we used single-reward bundles to compare neuronal response changes with behavioral changes. We established vector plots that displayed the ratio of reward weights (β’s) for z-scored neuronal population responses (Eq. **3**; Fig. 7*E-I*, red) and, separately, for behavioral choice (Eq. **1a**; green). The inequality of subjective value between the two rewards was manifested as deviation of these vectors from the 45 deg diagonal line. On-going reward consumption increased the elevation angle of the behavioral vector, indicating subjective value loss for Reward B (grape juice, water or mango juice) relative to Reward A (blackcurrant juice). The neuronal vector changed correspondingly (Fig. 7*E-I*, red). For example, during choice of the bundle (blackcurrant juice, grape juice) over zero-reward bundle, the behavioral vector angle increased from 40 deg before satiety to 65 deg during satiety, and the neuronal population vector increased from 35 deg to 62 deg (Fig. 7*E*, green, red). Similarly, during choice between two non-zero bundles, the behavioral vector increased from 40 deg to 52 deg while the neuronal vector increased from 38 deg to 45 deg (Fig. 7*F*). Further, the shorter neuronal vectors during satiety indicated general reduced responding, indicating additional general satiety (red). Bundles containing water or mango juice showed similar changes (Fig. 7*G-I*). Thus, both before and during satiety, the neuronal vectors (red) were within the CIs of the behavioral vectors (green), indicating intact neuronal subjective value coding that reflected the value changes with on-going reward consumption.

In addition to the vector analysis, IC slopes confirmed the close neuronal-behavioral correspondence during satiety, with satiety being defined by IPs exceeding the initial, pre-sated psychophysical CIs (Fig. S1*A, E*). As estimated from regression coefficient ratios (−β_2_ / β_1_) (Eq. **3**) and (−b / a) (Eq. **1**), the slopes of the linear neuronal ICs of single-reward bundles correlated with the slopes of linear behavioral ICs (Fig. 7*J*). These results from single-reward bundles during on-going reward consumption compared well with the results from the earlier OFC study on single rewards with spontaneously varying subjective reward value (1).

Taken together, the consumption-induced changes in single OFC neurons and in the OFC population corresponded well between single-reward and two-reward choice options. This similarity provides a sound foundation for the current findings on the more natural multi-component choice options.

## Discussion

This study investigated whether the reward responses of OFC neurons reflected reductions of subjective economic reward value induced by satiety for specific rewards, using stringent tests based on formal concepts of economic decision theory. As reward-specific satiety constitutes a major contributor to subjective value, the observed changes present crucial support in favor of economic value coding in OFC neurons and extend the role of this cortical structure in economic decision mechanisms. The use of bundles containing two differently sated rewards assured choice of both options as essential requirement for assessing subjective reward value in a controlled and unequivocal manner at choice indifference. The value loss was captured by graphic ICs that were constructed from psychophysically estimated IPs and represented the subjective economic value of the bundles. The ICs changed in an orderly and characteristic manner with on-going reward consumption (Figs. 1, 2, S1); they flattened progressively and changed from convex via near-linear to concave, thus indicating gradual subjective value loss for all bundle rewards (plotted on the x-axis) relative to blackcurrant juice (y-axis) (except for lesser peach juice value loss). These IC changes suggested that the animal required increasing quantities of the less sated blackcurrant juice for compensating the increasing subjective value loss of the other, more sated bundle reward with on-going consumption of both rewards. The specific and asymmetric IC changes made alternative explanations unlikely, such as general satiety, motivation loss, passage of time, or proximity of return to home cage, all of which would affect all rewards non-differentially. Licking behavior and consumption supported the notion of reward-specific satiety in a mechanism-independent manner (Figs. 3, S2, S3). Taken together, the demonstrated sensitivity to reward-specific satiety contributes an important argument in favor of formal subjective economic value coding by OFC neurons inferred from other choice data.

Our preceding study had established neuronal chosen value responses in OFC that were sensitive to multiple rewards and followed the animal’s rational choice of two-reward bundles, including completeness, transitivity and independence from option set size (24). The current study tested the effects of on-going reward consumption on these chosen value responses. We found that the neuronal responses matched the consumption-induced IC changes during recording periods of individual neurons. The responses became weaker for more sated rewards (Figs. 4, 5, S4, S5). Most impressively, neuronal responses failed to distinguish between bundles that were physically unchanged but came to be on the same ICs because of the ICs’ curvature change to concave (Figs. 4*D*). Classifiers predicting bundle discrimination from neuronal responses confirmed accurate subjective reward value coding both before and during satiety and demonstrated the substantial nature of neuronal changes (Figs. 6, S7–S9). Neuronal response vectors of conventional single-reward choice options correlated well with behavioral choice vectors; the correlations were maintained during consumption-induced subjective value changes (Fig. 7). These results demonstrate that OFC responses followed the differential subjective value changes induced by on-going reward consumption. As the rewards did not change physically with satiety, these results satisfy a necessary requirement for the coding of subjective value by OFC neurons that had been suggested based on choices between single-reward options (1) and two-reward bundles (30).

The consumption increase of sated rewards like water, grape juice, apple juice and mango juice (Figs. 3*H*, S2*A-D*) seemed to contradict earlier findings and the general intuition that satiety would rather reduce consumption of rewards on which an animal is sated. Differences in study design might explain these discrepancies. In the earlier studies, monkeys chose between sated and non-sated rewards, or between accepting and not accepting sated rewards, and naturally preferred non-sated rewards (8, 9, 15–17). As this tendency precluded choice indifference, we studied choice between bundles that each contained two differently sated rewards. Advancing satiety on Reward B required increasing quantities of this reward for choice indifference against bundles containing the less sated Reward A (see Figs. 1E, S1*B*). Thus, satiety increased consumption of the sated reward. By contrast, outright rejection of the more sated Reward B would have been represented by an upward sloped IC, which had been observed with lemon juice, yoghourt and saline (24) but not with the currently used rewards; such upward sloped ICs indicated that an animal needed to be ‘bribed’ with more reward for accepting these normally rejected sated rewards. By contrast, in the current study, the maintained downward IC slope indicated that the animal was not entirely averse to the sated reward.

The currently reported altered neuronal value coding with reward-specific satiety builds on previous studies on monkey OFC neurons that investigated satiety by clinical tests and behavioral observations. There, monkeys were presented with syringes or tubes containing various fruit juices; rating scales and go-nogo lick responses indicated behavioral acceptance or rejection of these juices (8, 9). The studies report that OFC neurons lost responses only for the particularly sated juice. Spontaneous variations of single-reward choices likely reflected appetite variations over the course of daily experimentation and affected chosen value coding in monkey OFC (1). Stronger general satiety effects in ventromedial prefrontal cortex compared to OFC (11) suggest widespread satiety sensitivity in ventral frontal cortex. Human studies using pleasantness ratings demonstrate neuroimaging correlates of reward satiety in OFC (12–14). The presence of satiety effects in OFC contrast with their absence in the earlier gustatory system, including nucleus of the solitary tract, frontal opercular taste cortex and insular taste cortex (6, 7, 31). The neurophysiological results (1, 8, 9, 11) correspond to the reward functions of frontal cortex that become deficient after lesion and inactivation, including associative strength (or, equivalently, incentive value), cognitive reward representation, and approach and goal-directed behavior (10, 15–17). Our present data on subjective value coding in a specific choice context extend these results into the domain of economic decision-making.

In addition to reward-specific satiety, on-going consumption induces also general satiety that may consist of a general loss of pleasure, arousal, attention and motivation. Our shorter neuronal population vectors with single-reward bundles may reflect general satiety, in addition to the changed vector angle that suggests reward-specific satiety (Fig. 7*E-I*, red). General and unspecific satiety would lead to general subjective value reduction in a similar way as reduced physical reward quantity; both would be represented by parallel, or curvi-parallel, IC displacement towards the x-y graph origin (Fig. 2*A-H*, left panels), whereas reward-specific satiety is indicated by changes in IC slope and curvature (right panels). General satiety is seen with reduced midbrain responses in humans during consumption of Swiss chocolate (12) and reduced dopamine responses in mice receiving food pellets for extended periods of time (32). Thus, general satiety cannot explain our asymmetric IC changes and the corresponding asymmetric neuronal changes, both of which reflect subjective value changes indicative of reward-specific satiety.

## Materials and Methods

The study used the same 2 male adult rhesus monkeys as previously (24, 30) and was licensed by the UK Home Office (for details, see Supplementary Information). The animals chose between two compound stimuli that were positioned at pseudorandomly alternating fixed left-right positions on a computer monitor and predicted bundles containing the same two rewards whose quantities varied pseudorandomly and without specific temporal order. We psychophysically estimated multiple choice indifference points (IP; Fig. 1*C*, S1*A, E*) to which we fitted ICs along which all bundles were equally preferred, using a hyperbolic function d:

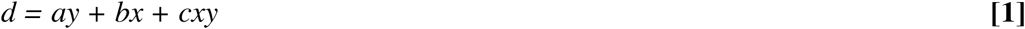

with *y* and *x* as milliliter quantity of Rewards A and B (Figs. 1*D, E*; S1*B, D, F*), *a* and *b* as weights of the influence of the two reward quantities, and *c* as curvature. Eq. **1** can be equivalently re-written as regression in analogy to the regression used for analysing neuronal responses:

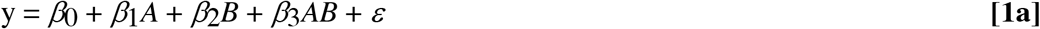

with *A* and *B* as milliliter quantity of Reward A (plotted at y-axis) and Reward B (x-axis), respectively, *β*_0_ as offset coefficient, *β*_1_ and *β*_2_ as behavioral regression coefficients, and *ε* as compound of errors *err*_0_, *err*_1_, *err*_2_, *err*_3_ for offset and regressors 1-3.

To test whether the animal’s choice reflected the quantity of the bundle rewards during satiety, rather than other, unintended variables such as spatial bias, we used the logistic regression:

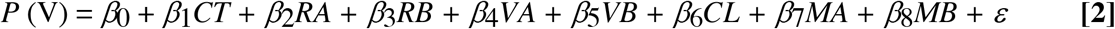

with *P* (V) as probability of choice of Variable Bundle, *β*_0_ as offset coefficient, *β*_1_ - *β*_7_ as correlation strength (regression slope) coefficients indicating the influence of the respective regressor, *CT* as trial number within blocks of consecutive trials, R*A* as quantity of Reward A of Reference Bundle, *RB* as quantity of Reward B of Reference Bundle, VA as quantity of Reward A of Variable Bundle, *VB* as quantity of Reward B of Variable Bundle, *CL* as choice of any bundle stimulus presented at the left, MA as consumed quantity of Reward A, *MB* as consumed quantity of Reward B, and *ε* as compound error for offset and all regressors.

Following behavioral training and surgical preparation for single neuron recording, we identified neuronal task relationships with the paired Wilcoxon-test. We identified changes of task-related neuronal responses across ICs with a linear regression:

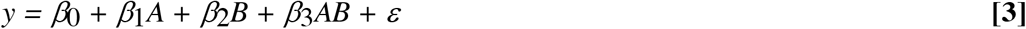

with *y* as neuronal response in any of the four task epochs (Bundle stimulus, Go, Choice, Reward), measured as impulses/s and z-scored normalized to the Pretrial control epoch of 1.0 s, *A* and *B* as milliliter quantity of Reward A (plotted at y-axis) and Reward B (x-axis), respectively, *β*_0_ as offset coefficient, *β*_1_ and *β*_2_ as neuronal regression coefficients, and *ε* as compound error. In addition, all significant neuronal response changes across ICs identified by Eq. **3** needed to be also significant in a Spearman rank-correlation test (*P* < 0.05).

To assess neuronal compliance with the two-dimensional IC scheme, we used a two-factor Anova on each task-related response that was significant for both regressors in Eq. **3**. Neuronal responses following the IC scheme were significant across-ICs (factor 1: *P* < 0.05) but insignificant within-IC (factor 2).

Chosen value (*CV*) was defined as:

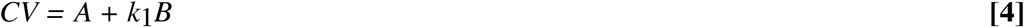

weighting parameter *k*_1_ served to adjust for differences in subjective value between rewards A and B, such that the quantity of Reward B entered the regression on a common-currency scale defined by Reward A. We assessed neuronal coding of chosen value in all neurons that followed the revealed preference scheme, using the following regression:

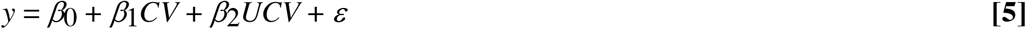

with U*CV* as value of the unchosen option that was not further considered here, and 𝝴 as compound error.

## Acknowledgements

We thank Aled David and Christina Thompson for animal and technical support, Polly Taylor for anesthesia during implantation, Paul Cisek for sharing his SQL-Matlab toolbox (NeuroMath), Charles R. Plott, Christopher Harris, Simone Ferrari-Toniolo and Fabian Grabenhorst for inspiration and insightful comments on experimental economics and neuronal data analysis. The Wellcome Trust (WT 095495, WT 204811), European Research Council (ERC; 293549) and US National Institutes of Mental Health Caltech Conte Center (NIMH; P50MH094258) supported this work.

## Supplementary information for

This Supplementary Information includes:

SI Methods
SI Figs. S1 to S9
SI Table S1

## SI Methods

### Animals

Two adult male macaque monkeys (*Macaca mulatta*; Monkey A, Monkey B), weighing 11.0 kg and 10.0 kg, respectively, were used in these experiments that had already yielded behavioral and neuronal data without satiety (1, 2). Neither animal had been used in any other study.

### Ethical approval

This research has been ethically reviewed, approved, regulated and supervised by the following institutions and individuals in the UK and at the University of Cambridge (UCam): the Minister of State at the UK Home Office, the Animals in Science Regulation Unit (ASRU) of the UK Home Office implementing the Animals (Scientific Procedures) Act 1986 with Amendment Regulations 2012, the UK Animals in Science Committee (ASC), the local UK Home Office Inspector, the UK National Centre for Replacement, Refinement and Reduction of Animal Experiments (NC3Rs), the UCam Animal Welfare and Ethical Review Body (AWERB), the UCam Governance and Strategy Committee, the Home Office Establishment License Holder of the UCam Biomedical Service (UBS), the UBS Director for Governance and Welfare, the UBS Named Information and Compliance Support Officer, the UBS Named Veterinary Surgeon (NVS), and the UBS Named Animal Care and Welfare Officer (NACWO).

### General behavior

The animals were habituated during several months to sit in a primate chair (Crist Instruments) for a few hours each working day. They were trained in a specific, computer-controlled behavioral task in which they contacted visual stimuli on a horizontally mounted touch-sensitive computer monitor (Elo) located 30 cm in front of them. The animal’s eye position in the horizontal and vertical plane were monitored with a non-invasive infrared oculometer (Iscan). Matlab software (Mathworks) running on a Microsoft Windows XP computer controlled the behavior and collected, analyzed and presented the data on-line. A solenoid valve (ASCO, SCB262C068) controlled by the same Windows computer served to deliver specific liquid quantities. A Microsoft SQL Server 2008 Database served for Matlab off-line data analysis. Following task training for about 6 months, animals were surgically implanted with a recording chamber for electrophysiological recordings, which typically lasted for another 6-10 months.

### Stimuli and rewards

A computer touch monitor presented the subject with two visual stimuli (4 deg apart), each representing one choice option, called Reference Bundle and Variable Bundle (Fig. 1*A*). Each bundle contained two rewards that were represented separately by a colored rectangle: Reward A, represented by a violet rectangle, and Reward B, represented by a green rectangle. Reward quantities were set independently and indicated by the vertical position of a bar within each rectangle (higher was more). The Reference Bundle contained two preset Reward quantities that were fixed for a given block of trials. The Variable Bundle contained a specifically set quantity of one Reward and an experimentally varied quantity of the other reward. Reward A in all bundles was blackcurrant juice without or with added monosodium glutamate (MSG), Reward B was grape juice, strawberry juice, mango juice, water, apple juice, peach juice, or grape juice with added inosine monophosphate (IMG). While MSG and IMG are known taste enhancers, we did not test choice preferences between them on their own. Thus, we cannot state whether they had own reward value for the animals.

### Task

Each trial began when the animal contacted a centrally located touch sensitive key for 1.0 s after a pseudorandom inter-trial interval of 1.6 ± 0.25 s. Then the two stimulus pairs representing the two bundles appeared at pseudorandomly alternating fixed left-right positions on a computer monitor in front of the animal (Fig. 1*B*). After 2.0 s, two blue spots appeared as GO stimulus underneath the bundle stimuli, upon which the animal released the touch key and touched the blue spot underneath the bundle of its choice within 2.0 s. The required action consisted of one arm movement and was constant across bundles and trials. After a hold time of 1.0 s, the blue spot underneath the chosen bundle turned green, and the blue spot underneath the unchosen bundle disappeared. Simultaneously a white frame around the chosen bundle appeared as feedback for successful choice. The computer-controlled liquid solenoid valve delivered Reward A at 1.0 s after the choice, followed 0.5 s later by Reward B (except when using peach juice as Reward B; here the sequence was reversed: Reward B was delivered first, then 0.5 s later Reward A, blackcurrant juice). Task training was initially restricted to one bundle type and was extended to other bundle types only when satisfactory behavioral performance was obtained.

The longer delay for liquid B compared to liquid A likely generated asymmetric temporal discounting that affected the subjective value of each liquid. However, all delays were kept constant, which allowed the subjective value differences from different temporal discounting to be incorporated as a constant factor into the subjective value of each liquid. We choose this delay, rather than simultaneous delivery or pseudorandomly alternating single liquid delivery, to prevent more serious taste interactions between simultaneously delivered liquids and to avoid temporal uncertainty. Reaching for a target before appearance of the blue dots, or key release during required key touch or target-hold, were considered as errors and lead directly to the inter-trial interval without reward.

### Estimation of behavioral ICs

The behavioral method for obtaining an IP from stochastic choice has been presented in full detail (1, 2). With two bundle options, the animal chose between the pre-set Reference Bundle (left in Fig. 1*A*) and the Variable Bundle (right) in repeated trials. Thus, the constant Reference Bundle provided a stable reference against the changing bundle composition in the Variable Bundle. We set one reward in the Variable Bundle to one unit (> 0.1 ml) above the quantity of the same reward in the Reference Bundle, while pseudorandomly varying the quantity of the other reward of the Variable Bundle over the whole test range and in pseudorandom temporal order within each block of trials. The variation of the animal’s repeated choice with that single, pseudorandomly varying reward allowed us to construct a full psychophysical function and estimate an IP from Weibull fitting (point of subjective equivalence; *P* = 0.5 choice of each bundle).

As in our previous study (1), we used the Matlab function GLMFIT for psychophysical fitting. This function returns a number called ‘Deviance’ between 0 and infinity that can be used to compare fitting between Weibull and logit. The Deviance is the difference between the log-likelihood of the fitted model and the maximum possible log-likelihood. Lower values are better. The estimated Deviance for psychophysics for the first 5,000 trials and 2 monkeys was 1.0415 for the Weibull model and 1.6009 for the logit model, suggesting that the Weibull fitted the data better. Hence, we used Weibull fitting for all psychophysical fitting.

We obtained each IP from a total of 80 trials (2 left-right stimulus positions with 5 equally spaced reward quantities in 8 trials). To avoid known adaptations in OFC neurons (3–6), we always tested the full reward range of the experiment.

To obtain an IC, we fit a series of IPs with a hyperbolic function d using weighted least mean squares:

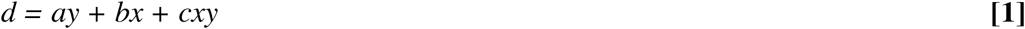

with y and x as milliliter quantity of Reward A (plotted on y-axis on 2D graph, Fig. 1*D*, *E*) and Reward B (plotted on x-axis), *a* and *b* as weights of the influence of the Reward quantities plotted on the y- and x-axes, respectively, and *c* as curvature. A potent reward that contributes strongly to the choice of the bundle would have a large weight (high coefficient a or b), whereas a less potent reward would have lower weight coefficients. Thus, with the potent (more weight) reward plotted on the x-axis, and the less potent (less weight) reward plotted on the y-axis, choice indifference between them would occur with smaller milliliter quantities on the x-axis compared to the y-axis. Hence, the IC slope would be steeper than the diagonal line (see Fig. 1*D, E*). By resolving Eq. **1** as *y* = −*(b / a) * x*, the IC slope would be the ratio of the coefficients that reflect the weights of the rewards: *−b / a*. With a higher potency of Reward B (x-axis) compared to Reward A (y-axis), the rectified IC slope would be larger than 1. Relatively stronger satiety for Reward B (x-axis) compared to Reward A (y-axis) would reduce the weight of Reward B, reduce the absolute value of the ratio *−b / a*, and flatten the IC slope. Thus, the IC slope *−b /* a describes the relative impact of the two bundle rewards (reflecting the value ratio between the two rewards), whereas the weights (*a* and *b*) describe the influence of the reward quantities.

The hyperbolic function can be re-written in an equivalent form to the regression with interaction used for analysing neuronal responses (see Eq. **3** below):

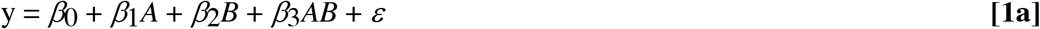

### Definition and criteria for pre-sated and sated states

With on-going reward consumption, the changes of psychophysical choice functions exceeding the confidence intervals (CI) of initial tests suggested a changed subjective value relationship between the two bundle rewards suggestive of relative, reward-specific satiety (see Figs. 1*D*, S1*A*, S1*E*). More specifically, the gradual effect of satiety on choice preference was identified by tracking the IPs as consumption advanced across blocks of 80 trials. Importantly, these changes occurred fast enough to be studied during the recording durations of single neurons, thus allowing us to compare responses between non-sated and sated states in the same neuron. The Weibull-fitted IPs were obtained psychophysically for fixed and equally spaced quantities of Reward B. Changes in relative subjective value of the two bundle rewards were assessed with interleaved anchor trials in choices between bundles with only one non-zero reward: bundle (fixed non-zero blackcurrant juice; no Reward B) vs. bundle (no blackcurrant juice; variable non-zero Reward B), using any Reward B (Fig. S1*B*). To aggregate IP data across sessions and compensate for across-session variability, we normalized the reward quantity ratio to the first titration block in all sessions. We then compared the normalized distributions of IPs within the CI of the first block with the distributions of IPs exceeding the CI of the first block.

### Control regressions for behavioral choice

To test whether the animal’s choice reflected the quantity of the bundle rewards during satiety, rather than other, unintended variables such as spatial bias, we used the logistic regression

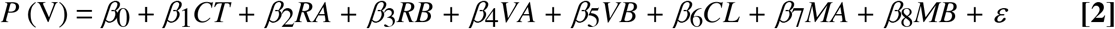

with *P* (V) as probability of choice of Variable Bundle, *β*_0_ as offset coefficient, *β*_1_ - *β*_7_ as correlation strength (regression slope) coefficients indicating the influence of the respective regressor, *CT* as trial number within blocks of consecutive trials, R*A* as quantity of Reward A of Reference Bundle, *RB* as quantity of Reward B of Reference Bundle, VA as quantity of Reward A of Variable Bundle, *VB* as quantity of Reward B of Variable Bundle, *CL* as choice of any bundle stimulus presented at the left, MA as consumed quantity of Reward A, *MB* as consumed quantity of Reward B, and *ε* as compound error for offset and all regressors. We used a binomial fit with logit link function to obtain standardized *β* coefficients. Choices over zero-reward bundles were excluded in the regression to avoid internal correlation between value and consumption.

### Licking

Licking was monitored with an infrared optosensor positioned below the juice spout (V6AP; STM Sensors). Anticipatory lick durations were measured between the appearance of the bundle stimuli and delivery of the first reward liquid (approximately 5 - 6 s duration) in bundles containing only one non-zero reward (single-reward bundles) within single working sessions. Licking data were collected with four bundle types, namely (blackcurrant juice, grape juice), (blackcurrant juice, water), (blackcurrant juice, strawberry juice) and (blackcurrant juice, mango juice).

### Surgical procedures and electrophysiology

As described before for the same animals (2), a head-restraining device and a recording chamber (40 x 40 mm, Gray Matter) were implanted on the skull under full general anesthesia and aseptic conditions. The stereotactic coordinates of the chamber enabled neuronal recordings of the orbitofrontal cortex (OFC) (7). We located the OFC from bone marks on coronal and sagittal radiographs taken with a guide cannula inserted at a known coordinate in reference to the implanted chamber, using a medio-lateral vertical and a 20º degree forward-directed approach aiming for area 13. Monkey A provided data from the left hemisphere and Monkey B from the right hemisphere via a craniotomy ranging from Anterior 30 to Anterior 38 and Lateral from 0 to 19. We conducted single-neuron electrophysiological recordings using both custom made glass-coated tungsten electrodes (8) and commercial electrodes (Alpha Omega) (impedance of about 1 MOhm at 1 kHz). Electrodes were inserted into the cortex with a multi-electrode drive (NaN drive) with the same angled approach as used for the radiography. Neuronal signals were collected at 20 kHz, amplified using conventional differential amplifiers (CED 1902 Cambridge Electronics Design) and band-passed filtered (high: 300 Hz, low: 5 kHz). We used a Schmitt-trigger to digitize the analog neuronal signal online into a computer-compatible TTL signal. However, we did not use the Schmitt-trigger to separate simultaneous recordings from multiple neurons, in which case we searched for another recording from only a single neuron, or we stored occasionally the data in analog form for off-line separation by dedicated software (Plexon offline sorter). An infrared eye tracking system monitored eye position (ETL200; ISCAN), with temperature check on an experimenter’s hand at the approximate position of the animal’s head.

### Definition of neurons following the revealed preference scheme

We analysed single-neuron activity during four task epochs vs. Pretrial control (1 s): visual Bundle stimulus (2 s), Go signal (1 s), Choice (1 s) and Reward (2 s, starting with Reward A, followed 0.5 s later by Reward B, except where noted, thus covering both rewards). To establish neuronal relationships to these task epochs, we compared the activity in each neuron during each task epoch separately against the Pretrial control epoch using the paired Wilcoxon test (*P* < 0.01). A neuron was considered task-related if its activity in at least one of the four task epochs differed significantly from the activity during the Pretrial control epoch.

Responses of individual neurons should follow the scheme of two-dimensional ICs that characterizes revealed behavioral preferences for two-dimensional bundles. Specifically, the responses should comply with three characteristics defined previously (2).

(Characteristic 1) Neuronal responses should change monotonically with increasing behavioral preference *across behavioral ICs*, irrespective to bundle composition. Such monotonic neuronal response changes should reflect increasing quantities of one or both bundle rewards, assuming a positive monotonic subjective value function on reward quantity.

(Characteristic 2) Neuronal responses should vary insignificantly for all equally preferred bundles positioned *along a same behavioral IC*, despite different physical bundle composition.

(Characteristic 3) Neuronal responses should follow the IC slope and the nonlinear curvature of behavioral ICs. The IC slope reflects the subjective value relationship between the two bundle rewards, and thus the subjective value of one reward (in our case Reward B) in the common currency of a reference reward (Reward A).

We used a combination of three statistical tests to assess these characteristics.

Characteristic 1: To capture the change *across ICs* in the most conservative, assumption-free manner possible, we used a simple linear regression on each Wilcoxon-identified task-related response:

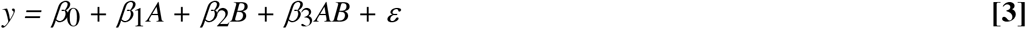

with *y* as neuronal response in any of the four task epochs, measured as impulses/s and z-scored normalized to the Pretrial control epoch of 1.0 s (z-scoring of neuronal responses applied to all regressions listed below), *A* and *B* as milliliter quantity of Reward A (plotted at y-axis) and Reward B (x-axis), respectively, *β*_0_ as offset coefficient, *β*_1_ and *β*_2_ as neuronal regression coefficients, and *ε* as compound error for offset and all regressors.

The coefficients *β*_1_ and *β*_2_ needed to be either both positive (indicating positive neuronal relationship, higher neuronal activity reflecting more reward quantity) or both negative (inverse neuronal relationship) to reflect the additive nature of the individual bundle components giving rise to revealed preference (*P* < 0.05, unless otherwise stated; t-test).

This linear regression assessed the degree of linear monotonicity of neuronal response change across ICs (*P* < 0.05 for *β* coefficients; t-test). Further, all significant positive or negative response changes identified by Eq. **3** needed to be also significant in a Spearman rank-correlation test that assessed ordinal monotonicity of response change across ICs without assuming linearity and numeric scale (*P* < 0.05).

Characteristics 1 and 2: To assess the two-dimensional *across/along IC* scheme in a direct and intuitive way, and without assuming monotonicity, linearity and numeric scale, we used a two-factor Anova on each Wilcoxon-identified task-related response that was also significant for both regressors in Eq. **3**; the factors were *across-IC* (ascending rank order of behavioral ICs) and *along-IC* (same rank order of behavioral IC). To be a candidate for following the IC scheme of Revealed Preference Theory, changes across-ICs should be significant (*P* < 0.05), changes within-IC should be insignificant, and their interaction should be insignificant.

Characteristic 3: Whereas the regression defined by Eq. **3** estimated neuronal responses across ICs, a full estimation of neuronal ICs for comparison with behavioral ICs would require inclusion of the IC slope and curvature, both of which depended on both rewards. By simplifying Eq. **3** by setting to zero both the *β*_3_ coefficient and the constant neuronal response along the IC, the neuronal IC slope would be the ratio of coefficients (−*β*_2_ / *β*_1_). Note the different meanings of the slope term: the neuronal IC slope (−*β*_2_ / *β*_1_) describes the relative coding strength of the two bundle rewards (reflecting the neuronal ratio of the two rewards), whereas each neuronal regression slope alone (*β*) describes the coding strength of neuronal response (correlation with the specific regressor). The neuronal IC curvature was estimated from the β_3_ coefficient of the interaction term AB (all *β*‘s *P* < 0.05; t-test).

### Neuronal chosen value coding

As stated before (2), chosen value (*CV*) was defined as the value of the option the animal was going to choose or had already chosen. As each option consisted of two components, we used a linear combination of the quantity of Reward A (blackcurrant juice) and Reward B (any of the other five rewards):

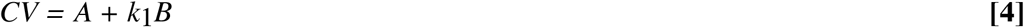

Weighting parameter *k*_1_ served to adjust for differences in subjective value between rewards A and B. We established parameter *k*_1_ during neuronal recording sessions from behavioral choice IPs using quantitative psychophysics in anchor trials (80 trials per test, see above Trial types for neuronal tests), rather than reading it from fitted ICs. Thus, *k*_1_ equals the ratio of coefficients *β*_2_ / *β*_1_ of Eq. 3.

We established a common-currency scale in ml for all tested rewards by defining blackcurrant juice or blackcurrant-MSG (Reward A) as reference (numeraire); the subjective value of any reward is expressed as real-number multiple k_1_ of the quantity of the numeraire at choice indifference. Specifically, the animal chose between the Variable Bundle that contained a psychophysically varied quantity of blackcurrant juice and the Reference Bundle that contained a fixed quantity of blackcurrant juice. A *k*_1_ of < 1 indicated that more quantity was required for choice indifference against blackcurrant juice; thus, *k*_1_ < 1 suggested that the tested reward had lower subjective value than blackcurrant juice. By contrast, *k*_1_ > 1 suggested higher subjective value, as less quantity was required for choice indifference.

We assessed the coding of chosen value and unchosen value in all neurons that followed the revealed preference scheme, using the following regression:

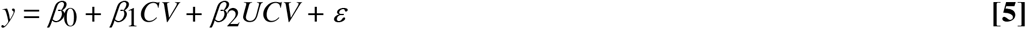

with *UCV* as value of the unchosen option that was not further considered here, and ε as compound error for offset and all regressors.

### Vector plots of OFC reward sensitivity

The purpose of this analysis was to provide quantitative and graphic information about satiety-induced behavioral and neuronal changes that would allow comparison with previous OFC studies that had not used two-component choice options with individually varying reward quantities and therefore did not establish ICs (9). This simplified analysis addressed monotonic response increase or decrease with increasing quantities of bundle rewards across ICs (characteristic 1 above), but did not address other IC characteristics such as trade-off, slope and curvature (characteristics 2 and 3) that had not been investigated previously. We established 2D plots whose dots indicated the relative contribution of each of the two bundle rewards to the neuronal response. We then compared vectors of behavioral choices with vectors of averaged neuronal population responses before and during satiety.

For behavioral choices, we plotted vectors (with 95% confidence intervals) from averaged dot positions defined by reward quantity (distance from center: sqrt (*β*_1_^2^ + *β*_2_^2^)) and relative weight (elevation angle: arctangent (*β*_1_ / *β*_2_)); coefficient *β*_1_ refers to Reward A (blackcurrant, y-axis), coefficient *β*_2_ refers to any of the other rewards (x-axis) (Eq. **1a**). The angle of the vector reflects the relative contribution the two bundle rewards to the choice, as estimated by the a and b coefficients (Eq. **1**). A deviation of the alignment angle from the diagonal line indicates an unequal contribution weight to bundle choice, and thus a non-1:1 reward ratio.

For neuronal responses, each dot on the two-dimensional plot was defined by the two β regression coefficients for neuronal responses (Eq. **3**; *P* < 0.01, t-test) for each of the two rewards in any of the four task epochs. The distance from center indicates the z-scored response magnitude (sqrt (*β*_1_^2^ + *β*_2_^2^)), coding sign (positive or negative), and relative weight (elevation angle; arctangent (*β*_1_ / *β*_2_)) of the two β coefficients. Coefficient *β*_1_ refers to Reward A (blackcurrant, y-axis), coefficient *β*_2_ refers to any of the other rewards (x-axis). Responses with negative (inverse) coding were rectified. Further IC characteristics such as systematic trade-off across multiple IPs and IC curvature played no role in these graphs. The alignment of the dots along the diagonal axis shows the relative coding strength for the two bundle rewards, as estimated by the β regression coefficients; a deviation from the diagonal line indicates an unequal influence of the two bundle rewards on the neuronal responses, reflecting a neuronal correlate of reward ratio.

### Neuronal decoding

We used a linear support vector machine (SVM) classifier to decode neuronal activity according to bundles presented at different behavioral ICs during choice over zero-reward bundle (bundle distinction) and, separately, according to the behavioral choice between two non-zero bundles located on different ICs (choice prediction). As in our main study on revealed preferences (2), we implemented the decoder with linear kernel using custom-written software with svmtrain and svmclassify procedures in Matlab R2015b (Mathworks). (our previous work had shown that use of nonlinear SVM kernels did not improve decoding) (10). The SVM decoder was trained to find the optimal linear hyperplane for the best separation between two neuronal populations relative to lower vs. higher ICs.

All analyses employed single-neuron data, consisting of single-trial impulse counts that had been z-normalised to the activity during the Pretrial epoch in all trials recorded with the neuron under study. The analysis included activity from all neurons whose responses followed the IC scheme of revealed preferences during any of the four task epochs, as identified by our three-test statistics, except where noted. The neurons were recorded one at a time; therefore, the analysis concerned aggregated pseudo-populations of neuronal responses.

The decoding analysis used 10 trials per neuron for each of two ICs (total of 20 trials). Extensive analysis suggested that higher inclusion of 15-20 trials per group did not provide significantly better decoding rates (while reducing the number of included neurons). For neurons that had been recorded with > 10 trials per IC, we selected randomly 10 trials from each neuron for each of the two ICs. We used a leave-one-out cross-validation method in which we removed one of the 20 trials and trained the SVM decoder on the remaining 19 trials. We then used the SVM decoder to assess whether it accurately detected the IC of the left-out trial. We repeated this procedure 20 times, every time leaving out another one of the 20 trials. These 20 repetitions resulted in a percentage of accurate decoding (% out of *n* = 20). The final percentage estimate of accurate decoding resulted from averaging the results from 150 iterations of this 20-trial random selection procedure. To distinguish from chance decoding, we randomly shuffled the assignment of neuronal responses to the tested ICs, which should result in chance decoding (accuracy of 50% correct). A significant decoding with the real, non-shuffled data would be expressed as statistically significant difference against the shuffled data (*P* < 0.01; Wilcoxon rank-sum test).

---

**Fig. S1.**
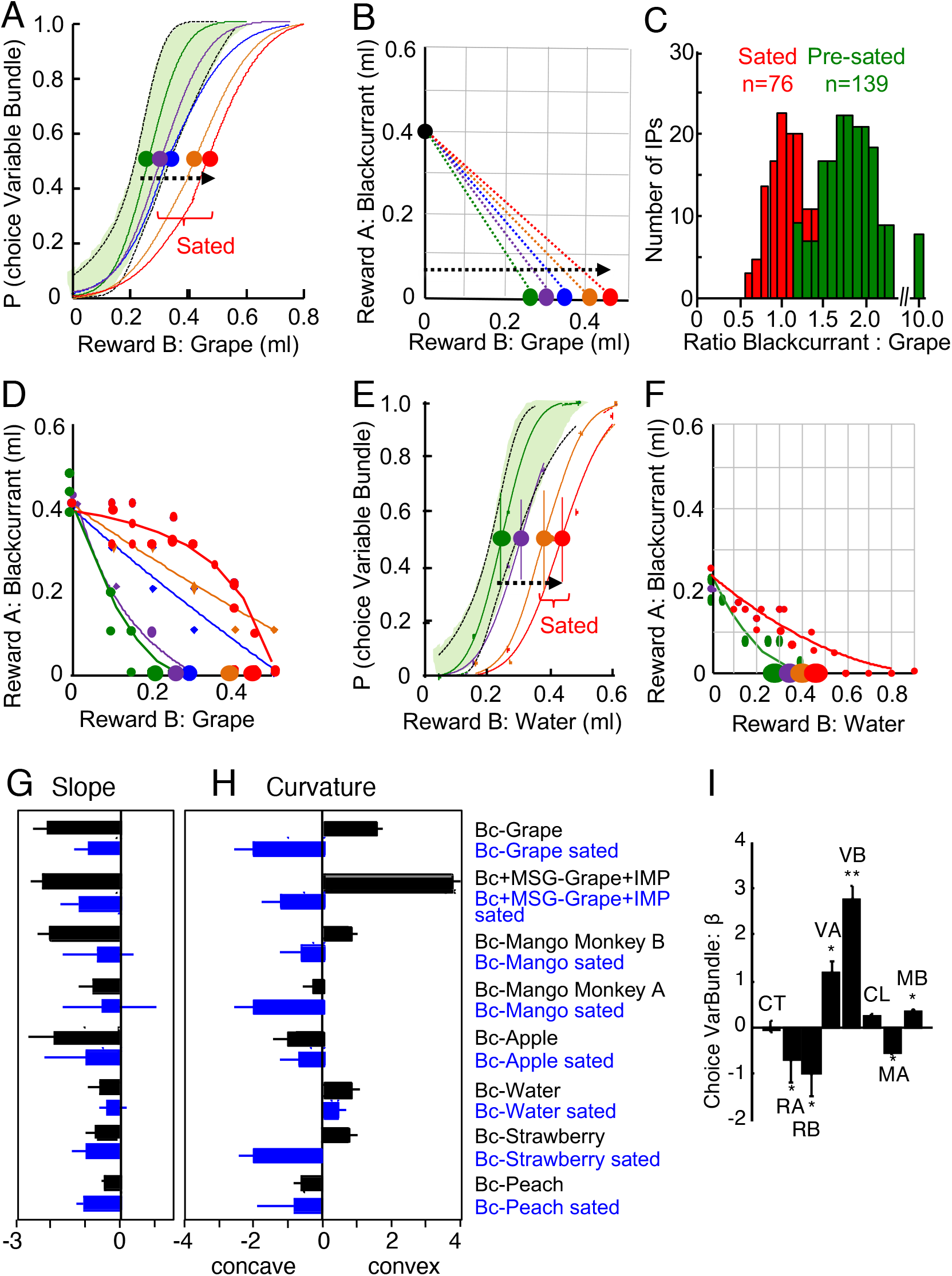
Additional behavioral tests demonstrating reward-specific satiety by changes of indifference curves (IC). (*A*) Psychophysical assessment of choice between single-reward bundles with grape juice variation (constant Reference Bundle: 0.4 ml blackcurrant juice, 0.0 ml grape juice; Variable bundle: 0.0 ml blackcurrant juice, varying grape juice). Green and violet curves inside green 95% confidence interval: initial choices; blue, orange and red curves: advancing consumption. The decrease in blackcurrant : grape juice ratio at IP was significant between the first IP and all IPs exceeding the first confidence interval (ratios of 1.9857 ± 0.0173, *n* = 139, green, vs. 1.0077 ± 0.02, orange and red; mean ± standard error of the mean, SEM; individual trial blocks: *P* = 9.6943 x 10^7^, Kolmogorov-Smirnov test; *P* = 2.336 x 10^−32^, Wilcoxon rank-sum test; *P* = 3.1712 x 10^−46^, t-test; Monkey A). Each curve and indifference point (IP) were estimated from 80 trials in a single block (Weibull fits). (*B*) Gradually developing relative satiety for grape juice indicated by increasing choice indifference points (IP; same bundles and animal as (A)): with on-going consumption of both juices, the animal required progressively more grape juice for choice indifference against the constant Reference Bundle (from green to red). The ratio blackcurrant : grape juice quantities at IP decreased from approximately 2:1 (0.4 ml of blackcurrant juice for 0.25 ml of grape juice, black vs. green dots) to about 1:1 (0.4 ml blackcurrant for 0.45 ml grape juice, black vs. red), suggesting subjective value loss of grape juice relative to blackcurrant juice. (*C*) Significant decrease of blackcurrant : grape juice ratio at IP with on-going consumption (same bundles as in A; Wilcoxon test). *n* = 139 and 76 IPs estimated in 43 trial blocks (Monkey A). (*D*) Choice tests between two-reward Variable bundle and Reference Bundle. Gradual change with grape juice indicate slope and curvature variation of ICs between pre-satiety (green, violet) and satiety (blue, orange, red) (single session; *n* = 2,960 trials; 80 trials/IP; Monkey A). (*E*), (*F*) Psychophysical tests and consumption-dependent IC change in Monkey B (constant Reference Bundle: 0.25 ml blackcurrant juice, 0.0 ml water; Variable bundle: 0.0 ml blackcurrant juice, varying water). With on-going consumption of both liquids, the animal gave up progressively more water for obtaining the same 0.25 ml of blackcurrant juice (from green to red), suggesting subjective value loss of water relative to blackcurrant juice. Same conventions as (*D*) (*n* = 2,400 trials; 80 trials/IP). (*G*), (*H*) Significant IC slope and curvature changes between pre-sated and sated states with on-going consumption of individual bundles, using regular test scheme (Fig. 1*E*) on standard two-reward bundles (Bc, blackcurrant juice; MSG, monosodium glutamate; IMP, inosine monophosphate; *P* = 0.0156 and *P* = 0.0313, respectively; Wilcoxon test). The slope parameter reflected the quantity ratio blackcurrant : other liquids at IP. (*I*) Value control by logistic regression for choice between Variable Bundle and Reference Bundle during satiety (Eq. **2**). According to significant *β* regression coefficients, choice of the Variable Bundle (Choice VarBundle) correlated significantly with quantity of rewards A and B in the Variable Bundle (VA, VB) and the Reference Bundle (RA, RB) and the consumed quantity of bundle rewards A (blackcurrant; MA) and B (various other liquids; MA). Choice varied insignificantly with consecutive trial number within blocks (CT) and left-right choice (CL). *n* = 7,243 trials pooled from several sessions; * *P* < 0.05; ** *P* < 0.01; t-test on *β*s.

**Fig. S2.**
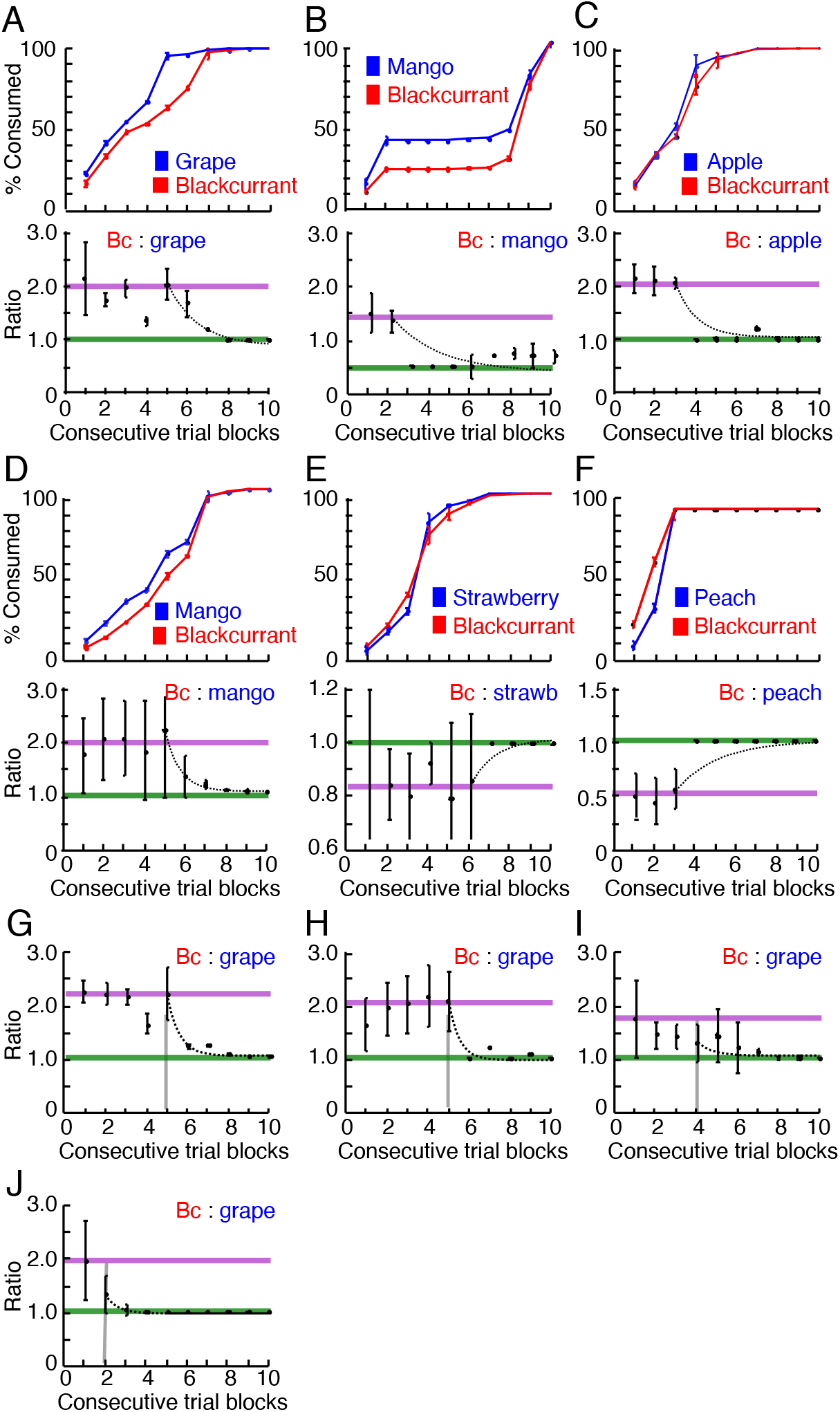
Consumption profiles and ratios of further test bundles indicating reward-specific satiety. (*A*) - (*F*) Cumulative juice consumption (top) and consumption ratios (bottom; pre-sated level: pink, sated level: green). Compared to blackcurrant juice (Bc; Reward B), on-going consumption increased significantly for the other bundle reward (Reward A) (*A*) grape juice: *P* = 1.364 x 10^−23^ (Monkey A; Kolmogorov-Smirnov test), (*B*) mango juice: *P* = 1.3566 x 10^−95^ (Monkey A); (*C*) apple juice: *P* = 0.045 (Monkey A); (*D*) mango juice: *P* = 3.2564 x 10^−22^ (Monkey B), changed insignificantly for (*E*) strawberry juice: *P* = 0.865 (Monkey A), and decreased significantly for (*F*) peach juice: *P* = 0.035 (Monkey B), using all trials (including bundles with two non-zero quantities). Trial numbers: (*A*): 9,812; (*B*): 5,200; (*C*): 2,160; (*D*): 12,570; (*E*): 8,201; (*F*): 440. Consumption ratio was measured only in anchor trials (each bundle contained only one liquid; see Fig. *S1A, B* for test scheme). Same conventions as Fig. 3*H, I*. (*G*) - (*J*) Stable consumption ratios for bundle (blackcurrant juice, grape juice) during one week (from Tuesday (*G*), to Friday (*J*); Monkey A). Note more rapid ratio change on Friday (*J*).

**Fig. S3.**
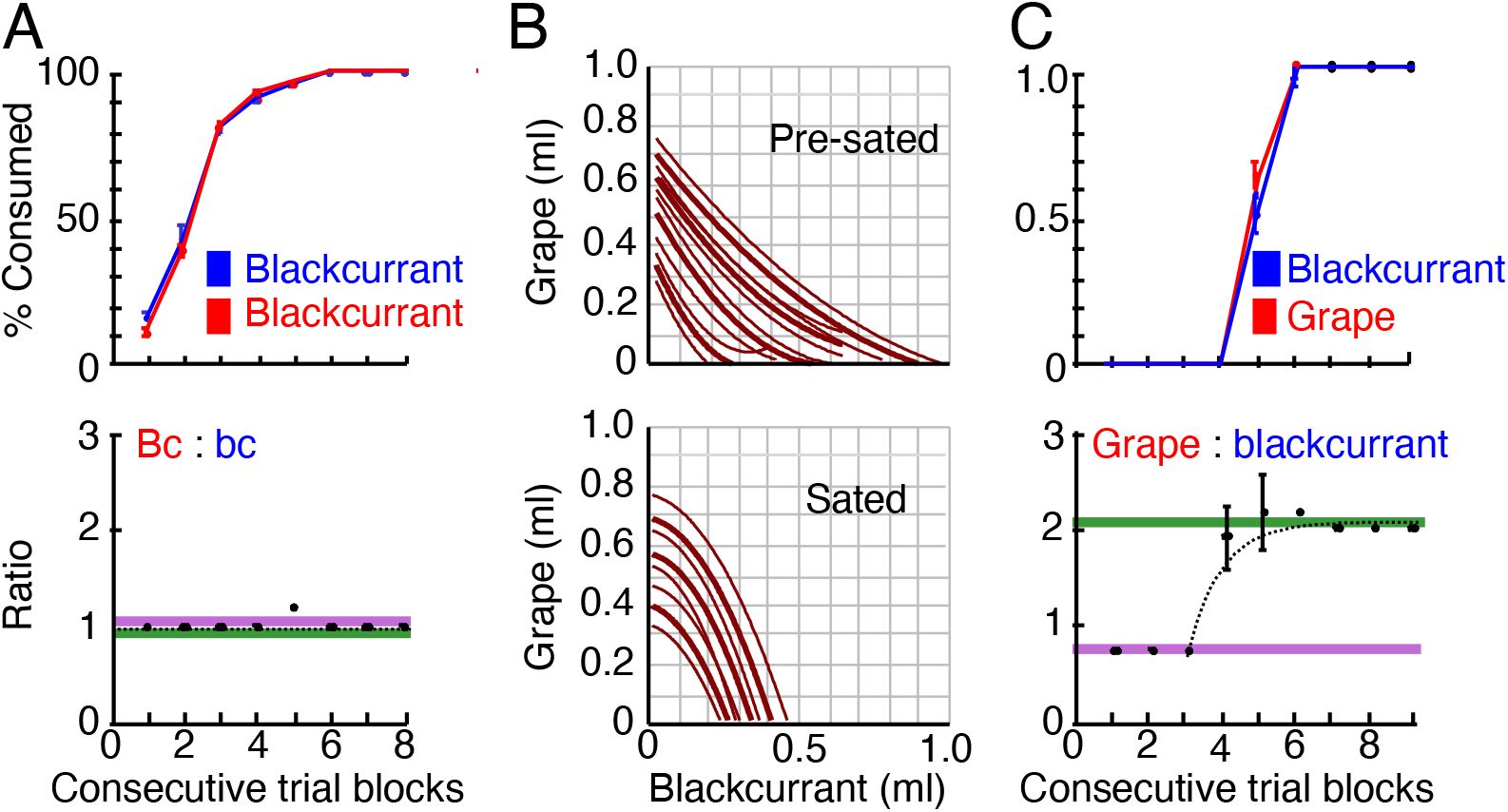
Reward-specific satiety is independent of juice delivery sequence. (*A*) Consumption profile and ratios for bundle containing blackcurrant (Bc) as both Reward A and Reward B: similar consumption and stable consumption ratio of both rewards. Pre-sated level: pink, sated level: green. For conventions, see Fig. S2. (*B*) Indifference curves (IC) for reversed delivery sequence of grape juice (first) and blackcurrant juice (second). Bottom: consistently concave ICs suggest stronger satiety for grape juice relative to blackcurrant juice: the animal required less blackcurrant juice for giving up the same quantity of grape juice after on-going consumption of both juices, indicating lower subjective value of grape juice and suggesting grape satiety. (*C*) Consumption profile and ratio for bundle (grape juice, blackcurrant juice) with opposite delivery sequence than usual: grape juice was delivered before blackcurrant juice.

**Fig. S4.**
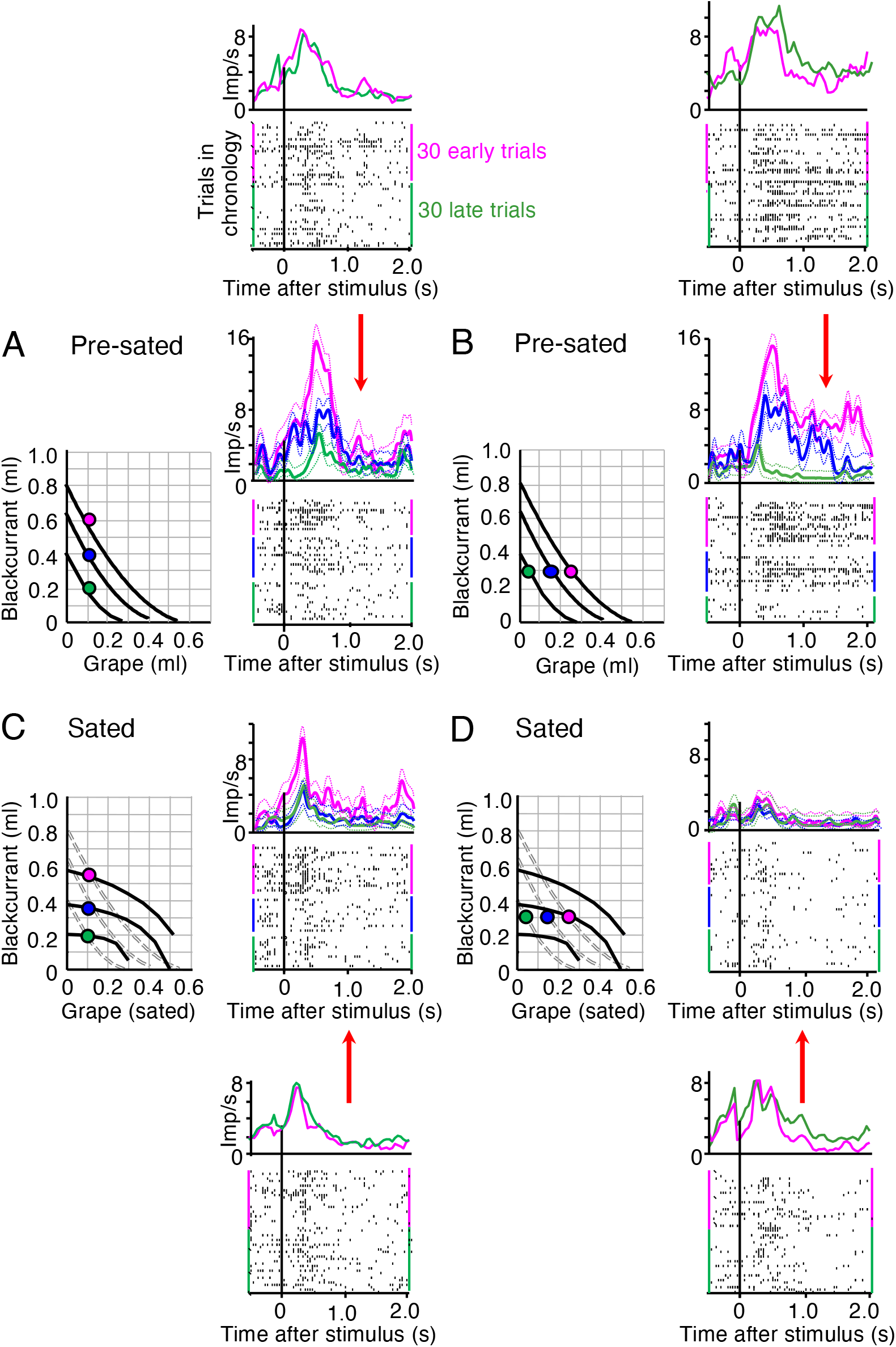
Satiety-related response change in single, chosen value coding OFC neuron is unrelated to trial sequence. Separate rasters and peri-stimulus time histograms in (*A*) - (*D*) show single-neuron discharges recorded in chronological order during early and late 30 trials of a session. The combined data are shown below and above the separate displays, as indicated by red arrows, together with their 95% confidence intervals. Same data and conventions as Fig. 4.

**Fig. S5.**
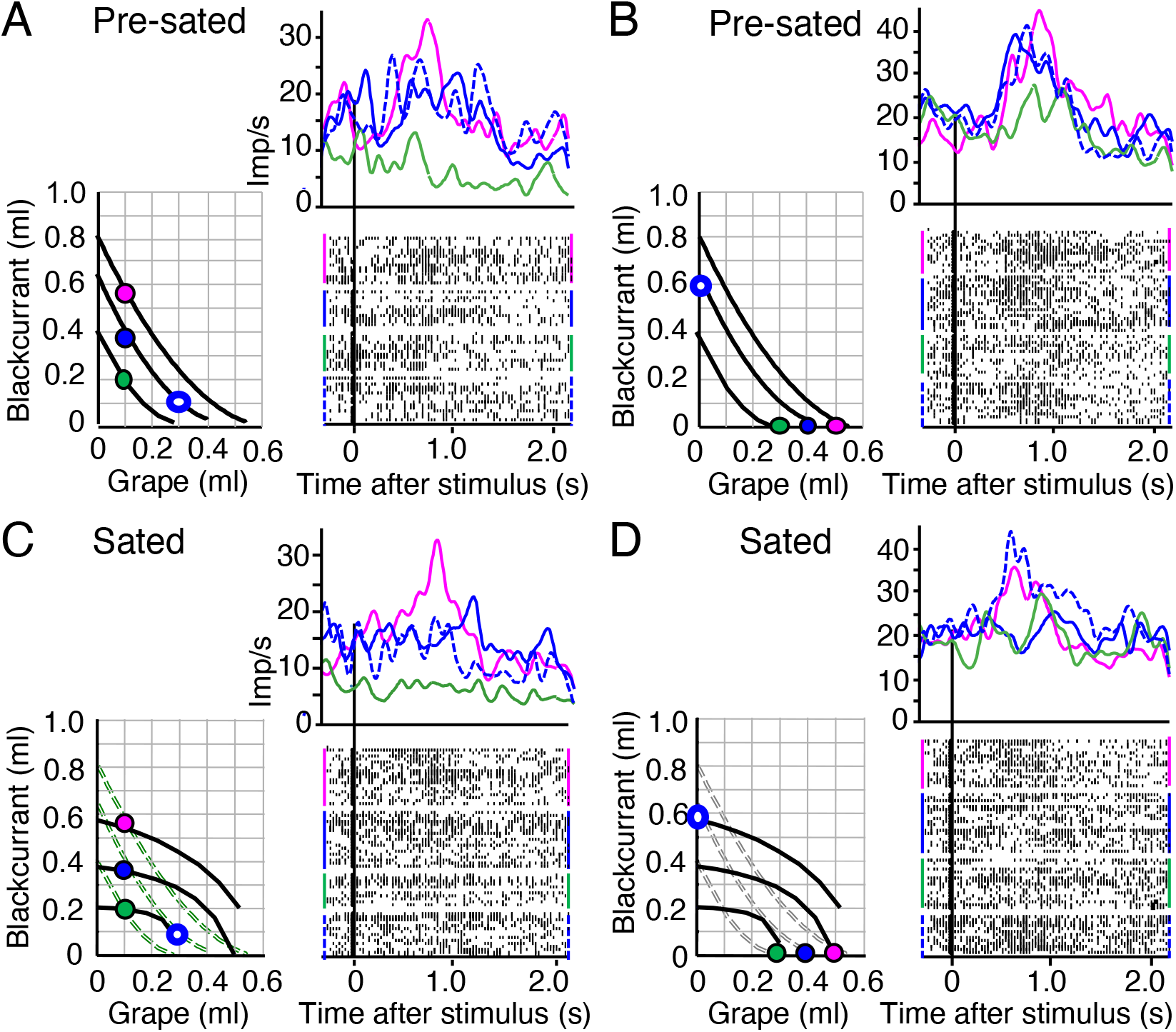
Satiety-related neuronal response change during choice between two non-zero bundles. (*A*) Significant monotonic neuronal response increase with value of chosen bundle across indifference curves (IC) before satiety (from green via solid blue to red) (*P* = 0.0015, F(1,64) = 10.12; *P* = 6.2183 x 10^−13^, F(2,64) = 58.0; two-way Anova: baseline vs. post-stimulus; across the 3 solid colored bundles). The animal chose between the Reference Bundle (hollow blue dot) and one of the Variable Bundles (solid colored dots). Responses to the two blue bundles on the same IC (representing equal preference) varied insignificantly despite different juice composition (*P* = 0.5488; t-test). Response to Reference Bundle (hollow blue dot) is indicated by dotted line. Binwidth 10 ms, Gaussian kernel smoothing. (*B*) As (*A*) but for grape juice variation. Responses varied significantly across ICs with grape juice (*P* = 5.0301 x 10^−5^, F(1,116) = 16.5; *P* = 1.0999 x 10^−9^, F(2,116) = 23.17). Responses to the two blue bundles on the same IC differed insignificantly (P = 0.2622). Same color labels as (*A*). (*C*) Despite IC change indicating satiety, the neuronal response increase across ICs remained significant (*P* = 2.3892 x 10^−5^, F(1,102) = 12.921; *P* = 4.87835 x 10^−12^, F(2,102) = 48.35). However, the two unchanged blue bundles were now on different ICs, and their responses correspondingly varied significantly (*P* = 0.0028). (*D*) With slope and curvature change indicating satiety, the three bundles with grape juice variation were now located within only two ICs. Although the neuronal response increase across ICs remained significant (*P* = 0.0071, F(1,106) = 4.97; *P* = 5.70 x 10^−7^, F(2,106) = 25.18), the peak response was reduced by 25% (from 40 to 30 imp/s, red) and the three responses were closer to each other. Further, the two unchanged blue bundles were now on different ICs, and their responses now differed significantly (*P* = 0.0201). Thus, the changes of neuronal responses were consistent with the IC change indicating satiety.

**Fig. S6.**
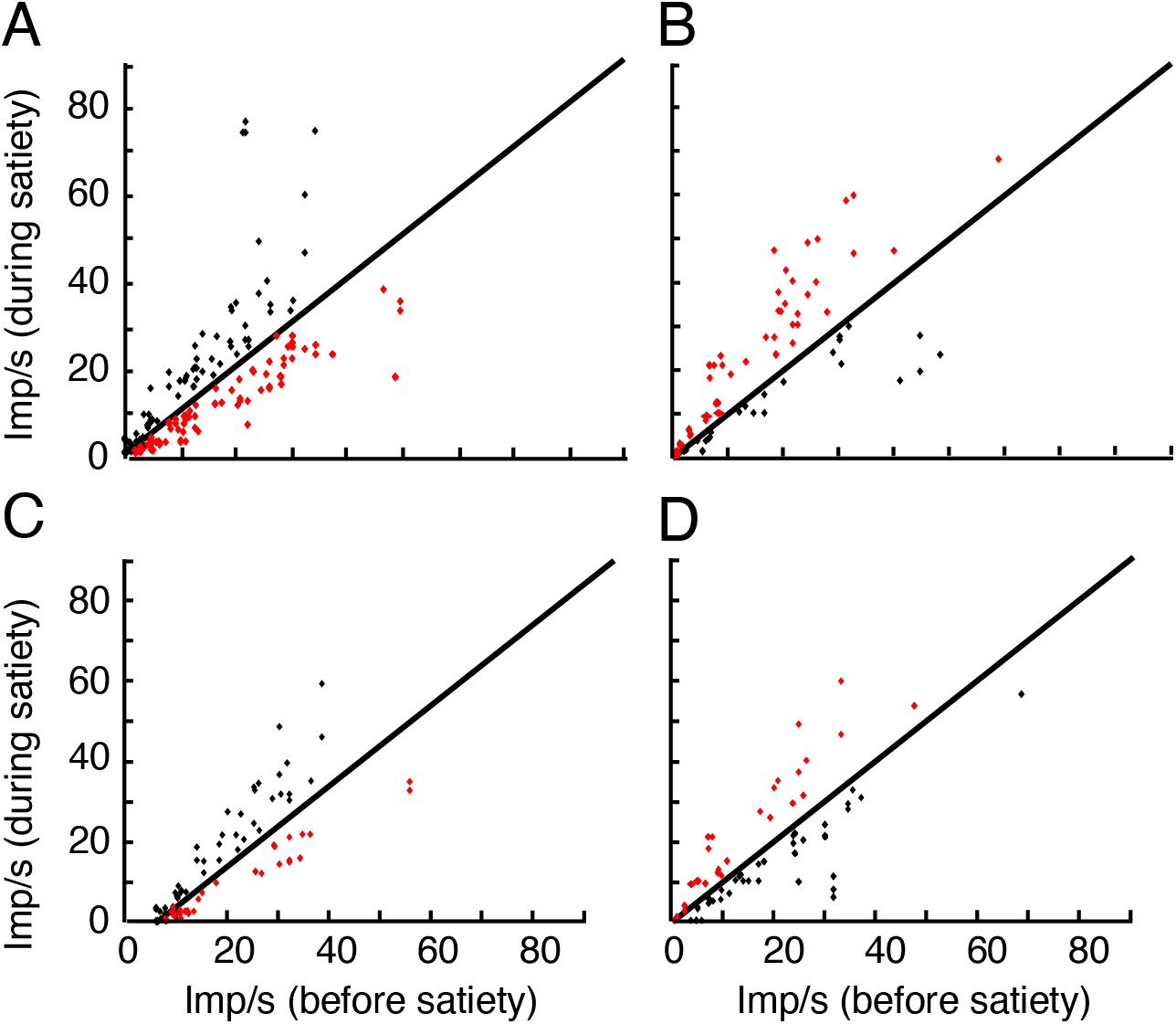
Numeric display of satiety-induced neuronal response changes. (*A*) Response changes in positive coding chosen value neurons in any of the four task epochs (Bundle stimulus, Go, Choice and Reward; Table S1) during choice over zero-reward bundle. Red: significant response decrease in population reflecting satiety-induced subjective value reduction (*P* = 7.15 x 10^−4^; 101 responses in 31 neurons; 1-tailed t-test). Black: significant response increase (*P* = 0.0014; 69 responses in 21 neurons). Imp/s: impulses/second. (*B*) As (*A*) but for negative (inverse) value coding neurons. Red: significant response increase reflecting satiety-induced subjective value reduction (*P* = 0.0013; 54 responses in 15 neurons). Black: insignificant response decrease (*P* = 0.1274; 33 responses in 14 neurons). (*C*) As (*A*) but for choice between two non-zero bundles. Red: response decrease (*P* = 0.0156; 54 responses in 16 neurons; 1-tailed t-test). Black: response increase (*P* = 0.0101; 57 responses in 16 neurons). Imp/s: impulses/second). (*D*) As (*B*) but for choice between two non-zero bundles. Red: significant response increase (*P* = 0.0242; 31 responses in 9 neurons). Black: insignificant response decrease (*P* = 0.1939; 36 responses in 14 neurons).

**Fig. S7.**
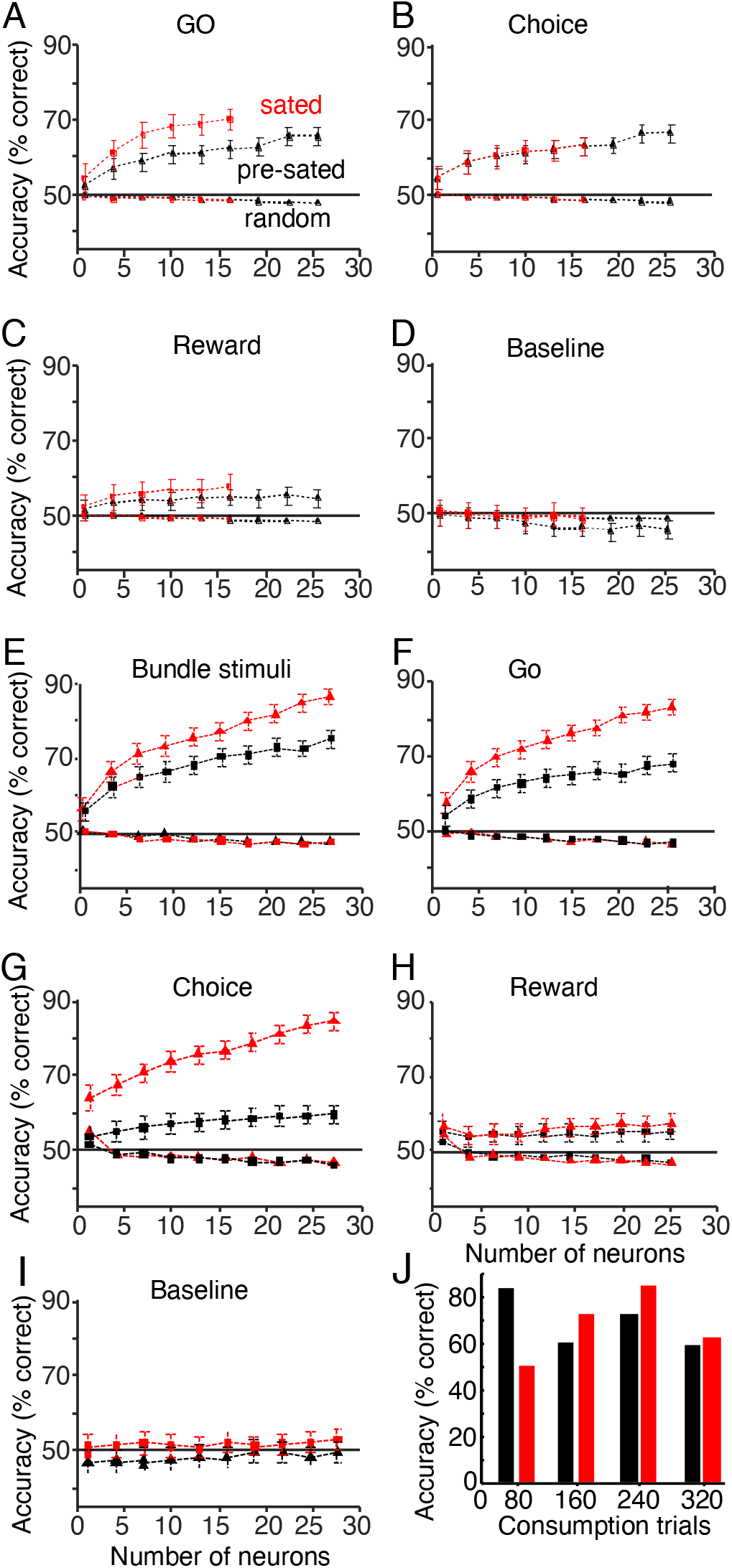
Satiety-induced changes in bundle classification during specific task epochs. (*A*) - (*D*) Bundle classification by support vector machine using neuronal chosen value responses to stimuli of bundles positioned on the lowest and third lowest indifference curve, respectively (choice over zero-reward bundle). The classifier was trained on neuronal responses collected during satiety and tested during satiety (red) and before satiety (black). Baseline refers to 1 s Pretrial control epoch before Bundle stimuli. Random: control classification with shuffled assignment of neuronal responses to ICs. For conventions, see Fig. 6. (*E*) - (*I*) As (*A*) - (*D*) but for choice prediction by neuronal responses during choice between two non-zero bundles. (*J*) Classification accuracy of neuronal responses with advancing liquid consumption. Same data selection as for (*A*) - (*D*)) and collapsed across all task epochs. Black: before satiety, red: during satiety.

**Fig. S8.**
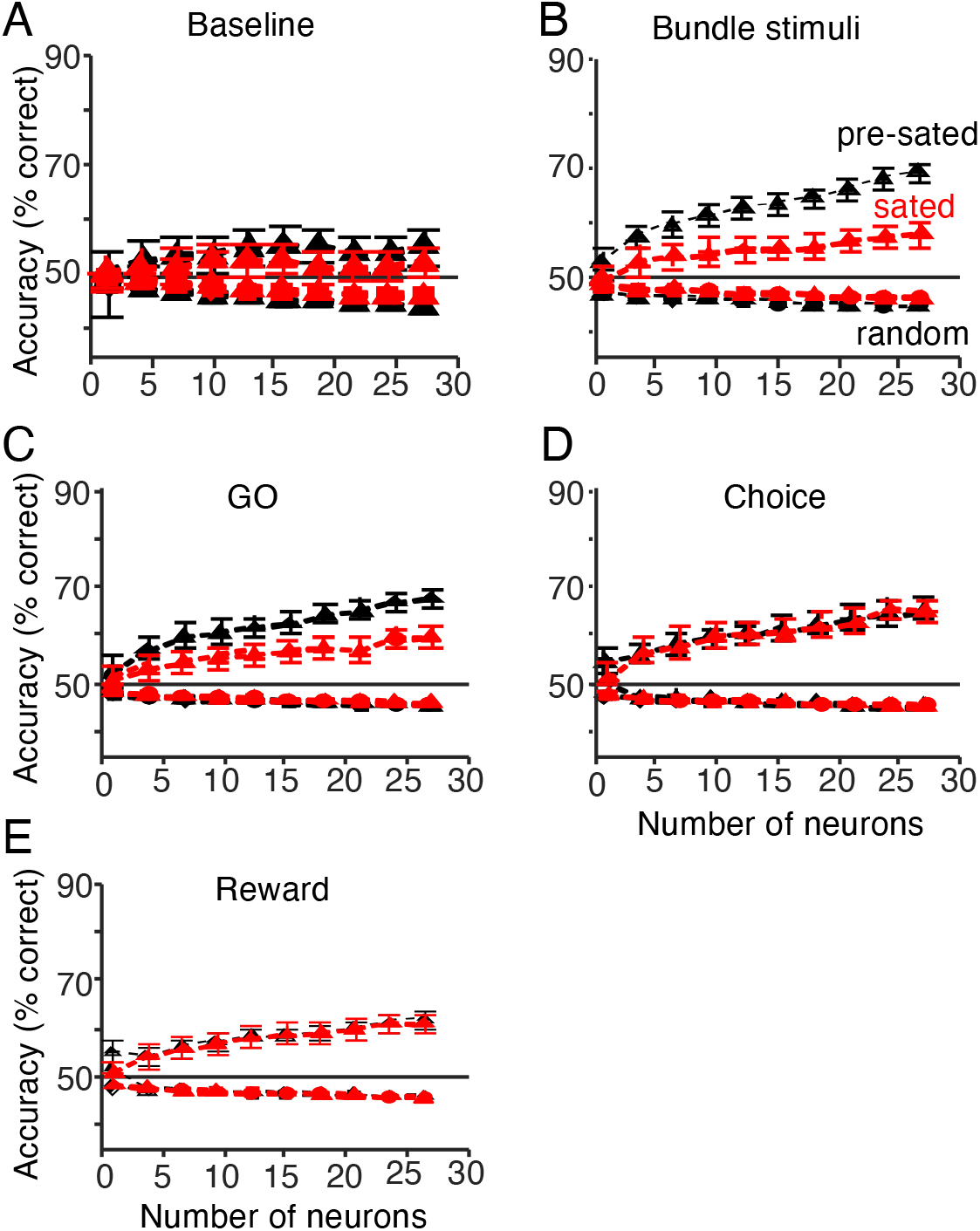
Satiety-induced changes in bundle classification derived from unmodulated and unselected neuronal responses. Prediction of bundle choice between lowest and third lowest indifference curves from activity of 265 unselected neurons during the five task epochs (choice between two non-zero bundles). The classifier was trained on neuronal responses before satiety and tested for bundle distinction before satiety (black) and during satiety (red). Random: control with shuffled assignment of neuronal responses to ICs. For conventions, see Figs. 6 and S7.

**Fig. S9.**
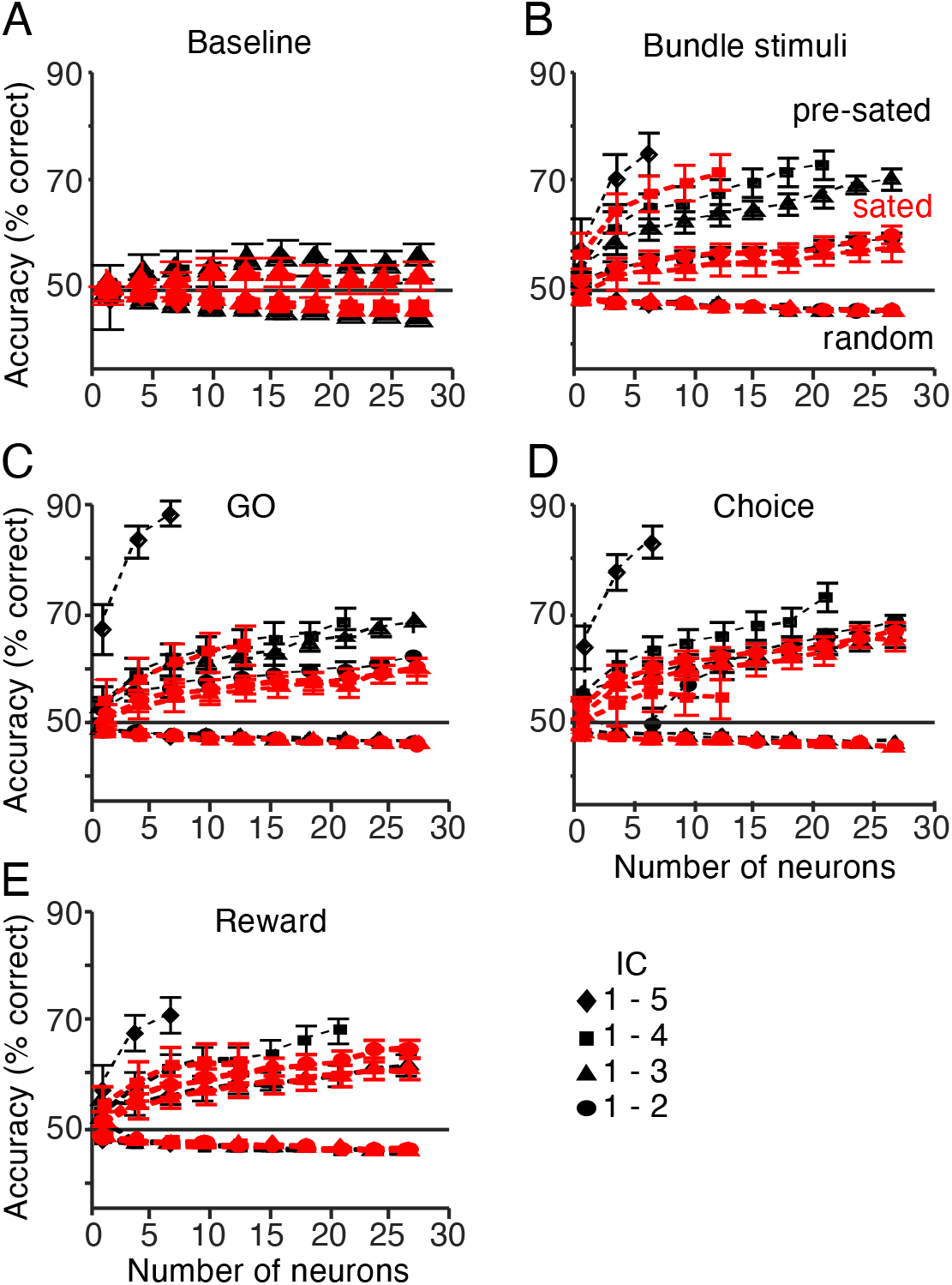
Satiety-induced changes in bundle classification derived from unmodulated and unselected neuronal responses: all possible comparisons. Prediction of bundle choice from activity of 265 unselected neurons during the five task epochs (choice between two non-zero bundles). Compared to Fig. S8, all possible binary comparisons between indifference curves (IC) were tested; ICs are numbered from bottom to top, and the four comparisons are indicated by symbols. For conventions, see Fig. S8.

**Table S1.** Neuronal changes with on-going reward consumption during different task epochs. The table shows data from chosen value responses, separated according to the four task epochs (Bundle stimulus, Go, Choice and Reward). Data are from all bundles tested for satiety (Reward A: blackcurrant juice, Reward B: grape juice, water or mango juice). Positive coding refers to response increase with higher subjective value before satiety, whereas negative coding refers to response decrease with higher subjective value. Most neurons were tested both in choice over zero-reward bundle and in choice between two non-zero bundles. Changes during the Reward epoch may indiscriminately reflect changes in subjective reward value and consumption (mouth movements, sensory stimulation); no attempts were made to distinguish neuronal relationships between these factors.

| Choice over zero-reward bundle |  |  |  |  |  |  |
| --- | --- | --- | --- | --- | --- | --- |
| Task epoch | Neurons tested | Neurons | Responses | Neurons | Responses | Neurons |
|  |  | Response decreases |  | Response increases |  | No effects |
| Positive coding |  |  |  |  |  |  |
| Bundle stimulus |  |  | 30 |  | 21 |  |
| Go |  |  | 28 |  | 17 |  |
| Choice |  |  | 25 |  | 15 |  |
| Reward |  |  | 18 |  | 16 |  |
| Subtotal | 55 | 31 | 101 | 21 | 69 | 3 |
| Negative coding |  |  |  |  |  |  |
| Bundle stimulus |  |  | 10 |  | 15 |  |
| Go |  |  | 8 |  | 15 |  |
| Choice |  |  | 8 |  | 11 |  |
| Reward |  |  | 7 |  | 13 |  |
| Subtotal | 31 | 14 | 33 | 15 | 54 | 2 |
| Choice between two non-zero bundles |  |  |  |  |  |  |
| Task epoch | Neurons tested | Neurons | Responses | Neurons | Responses | Neurons |
|  |  | Response decreases |  | Response increases |  | No effects |
| Positive coding |  |  |  |  |  |  |
| Bundle stimulus |  |  | 16 |  | 16 |  |
| Go |  |  | 15 |  | 15 |  |
| Choice |  |  | 13 |  | 16 |  |
| Reward |  |  | 10 |  | 10 |  |
| Subtotal | 38 | 16 | 54 | 16 | 57 | 6 |
| Negative coding |  |  |  |  |  |  |
| Bundle stimulus |  |  | 11 |  | 9 |  |
| Go |  |  | 9 |  | 8 |  |
| Choice |  |  | 8 |  | 6 |  |
| Reward |  |  | 8 |  | 8 |  |
| Subtotal | 32 | 14 | 36 | 9 | 31 | 9 |

